# Unraveling Oxidative Stress Resistance: Molecular Properties Govern Proteome Vulnerability

**DOI:** 10.1101/2020.03.09.983213

**Authors:** Roger L. Chang, Julian A. Stanley, Matthew C. Robinson, Joel W. Sher, Zhanwen Li, Yujia A. Chan, Ashton R. Omdahl, Ruddy Wattiez, Adam Godzik, Sabine Matallana-Surget

## Abstract

Oxidative stress alters cell viability, from microorganism irradiation sensitivity to human aging and neurodegeneration. Deleterious effects of protein carbonylation by reactive oxygen species (ROS) make understanding molecular properties determining ROS-susceptibility essential. The radiation-resistant bacterium *Deinococcus radiodurans* accumulates less carbonylation than sensitive organisms, making it a key model for deciphering properties governing oxidative stress resistance. We integrated shotgun redox proteomics, structural systems biology, and machine learning to resolve properties determining protein damage by γ-irradiation in *Escherichia coli* and *D. radiodurans* at multiple scales. Local accessibility, charge, and lysine enrichment accurately predict ROS-susceptibility. Lysine, methionine, and cysteine usage also contribute to ROS-resistance of the *D. radiodurans* proteome. Our model predicts proteome maintenance machinery and proteins protecting against ROS are more resistant in *D. radiodurans*. Our findings substantiate that protein-intrinsic protection impacts oxidative stress resistance, identifying causal molecular properties.

**One Sentence Summary:** Proteins differ in intrinsic susceptibility to oxidation, a mode of evolutionary adaptation for stress tolerance in bacteria.

## Main Text

Proteome oxidation caused by reactive oxygen species (ROS) is a primary determinant of cellular sensitivity to desiccation and irradiation (*1, 2*) and is involved in the progression of age-related human diseases, including neurodegeneration and cancer (*3*). ROS toxicity is a common antibiotic mechanism (*4*) and presents challenges in biotechnology including metabolic engineering (*5–7*) and synthetic systems involving the high expression of fluorescent proteins (*8*).

Prior to the previous decade, the dogma surrounding biological sensitivity to ionizing radiation focused primarily on DNA damage, but this changed as key experiments substantiated the role of protection from protein oxidation in the extreme radioresistance of the bacterium *Deinococcus radiodurans* (*9*). *Deinococcus* is an ideal model for investigating resistance to ROS because of its notorious tolerance of extreme oxidative stress, even prolonged cosmic doses of γ-radiation (*10*). *D. radiodurans* accumulates less protein oxidation than more sensitive species such as *Escherichia coli* (*2*). Resistance in *D. radiodurans* is due partly to highly active ROS-detoxifying systems providing protein-extrinsic protection against ROS (i.e. not a property of the oxidation targets themselves) (*1, 11*); however, the extent to which protein-intrinsic properties (i.e. specific to individual protein species) contribute to ROS resistance and how such properties are distributed across distinct protein species has not been well-studied. A comparative study of *radiodurans* and *E. coli* proteomes offers an opportunity to identify proteins with distinguished vulnerability to ROS, thereby discovering mechanisms that contribute to the survival of oxidative stress following irradiation.

ROS damage proteins by the oxidation of side chains and backbones generally resulting in loss of function due to misfolding, aggregation, and proteolysis. Protein carbonyl sites (CS) on arginine, lysine, proline, and threonine (RKPT) sidechains (Fig. S1) present the most severe oxidative damage due to their irreversibility and frequency of occurrence. Furthermore, these carbonyls themselves are also highly reactive leading subsequently to additional damaging downstream reactions, such as non-enzymatic backbone cleavage via the proline oxidation pathway (*12–14*). In this way, RKPT carbonylation can be thought of as a committed step initiating a cascade of protein damage. Site-specific susceptibility to oxidation differs across amino acid types and structural location, extending to the whole-molecule scale to distinguish ROS vulnerability across protein species (Fig. 1A). However, specific molecular properties responsible for this vulnerability remain poorly understood.

**Fig. 1.**
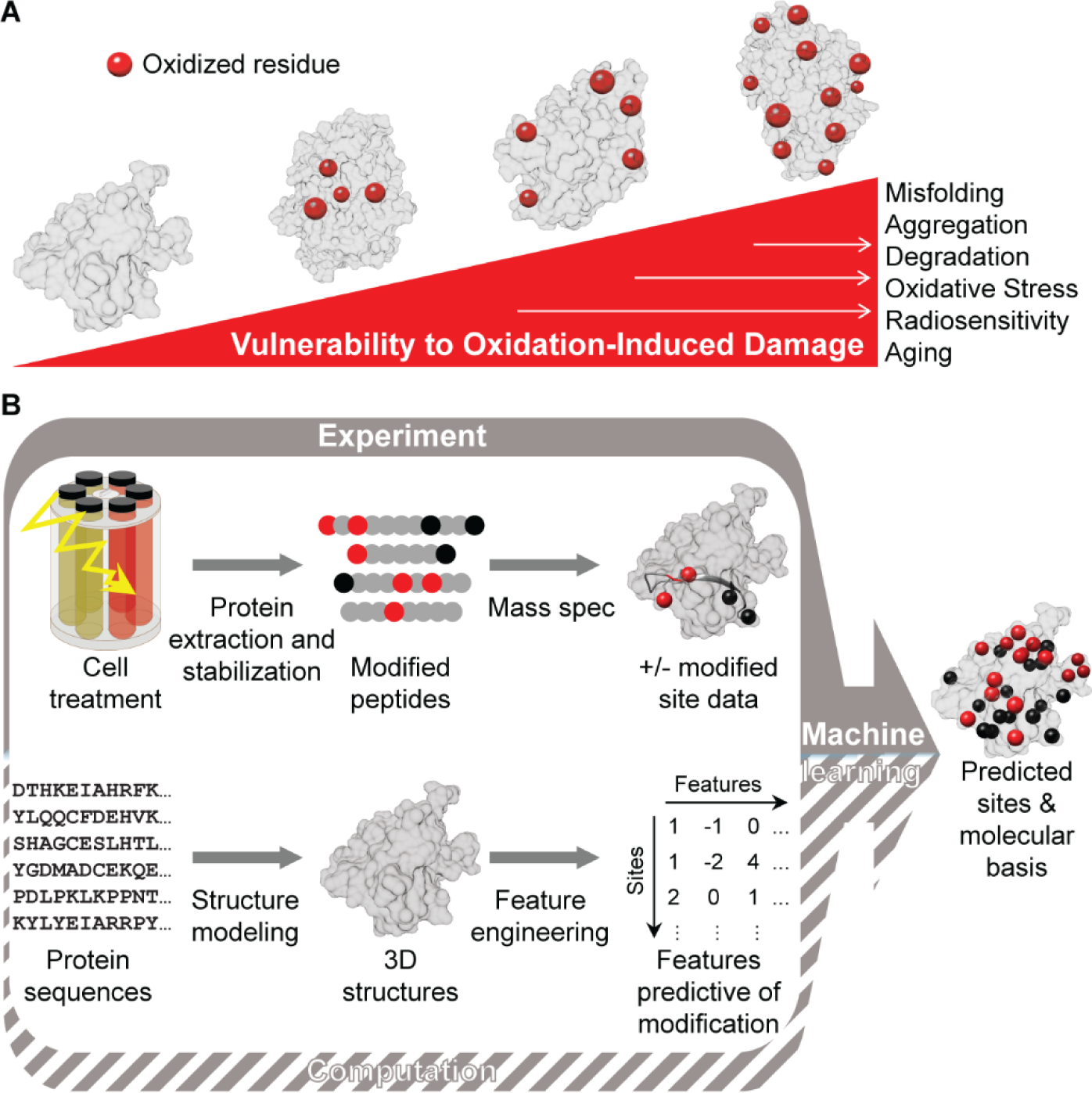
Study concept and workflow. (A) Relationship between oxidation site distribution, protein vulnerability to reactive oxygen species, and stress phenotypes. (B) Structural systems biology workflow for proteome-wide carbonyl site prediction. Red circles = carbonyl sites (CS); black circles = non-oxidized RKPT residues; gray protein regions = non-RKPT residues.

Previous work provided evidence that there is a difference in oxidation susceptibility between distinct protein species in bacteria through observation of banding patterns on carbonyl assay gels (*15*), but this work did not provide protein identification, quantification, nor residue-specificity of carbonylation events. Identification of proteins prone to oxidation and their specific oxidation sites is vital to understanding the molecular manifestation of deleterious oxidative stress phenotypes. This goal has motivated the development of mass spectrometry for direct proteome-wide CS identification and concomitant relative abundance changes, termed shotgun redox proteomics (*16*). However, these experiments provide limited coverage of modified sites, a common problem in proteomics of post-translational modifications. Chemical derivatization during these experiments helps to stabilize the inherently transient, highly reactive protein carbonyls to promote their detection, but interference from derivatized adducts with proteolytic sites can also limit CS sampling capabilities. Computational methods for carbonylation site prediction are intended to learn shared features across oxidation sites in redox proteomic datasets and generalize to unknown sites on other proteins. Existing methods (*17–19*) are not ideal because they rely on linear sequence motifs and local homology; such a correlative basis for predicting structure-function relationships can require a very large number of example sequences before very strong predictors can be trained (*20*), which are not yet available in the context of redox proteomic data. Furthermore, the exclusion of molecular structure features beyond simple sequence motifs provides limited understanding of causal mechanisms for protein carbonylation.

In response to the limitations of conventional techniques, the field of structural systems biology offers approaches based on protein 3D molecular properties to investigate multi-scale proteomic questions, including mechanisms of physicochemical stress (*21, 22*). These approaches are empowered by the expansion of experimentally-determined protein structures and advances in protein fold prediction (*23*). Our robust experimental design combined for the first time redox proteomics performed on cells exposed to an acute-dose of γ-radiation with structural systems biology and machine learning (Fig. 1B), generating a predictive model for protein carbonylation. This interdisciplinary workflow enabled proteome-wide characterization of susceptibility to oxidation in *E. coli* and *D. radiodurans*, identifying phenotypically-important protein targets, providing molecular explanations for target susceptibility, and broadly establishing the role of protein-intrinsic properties in the survival of extreme oxidative stress.

## Results and Discussion

### 1. Gamma-irradiation causes more targeted protein oxidation in *D. radiodurans* than *E. coli*

To investigate oxidative damage to bacterial proteins, cultures were exposed to an acute dose of γ-radiation lethal to *E. coli* but yielding 55-70% survival of *D. radiodurans*, and protein carbonyls and relative abundance changes were measured by mass spectrometry (Fig. 1B, Fig. 2, Fig. S2). In order to limit *de novo* protein synthesis throughout and following irradiation, bacterial cultures were maintained near 0°C using a custom rack design (Files S1 and S2).

**Fig. 2.**
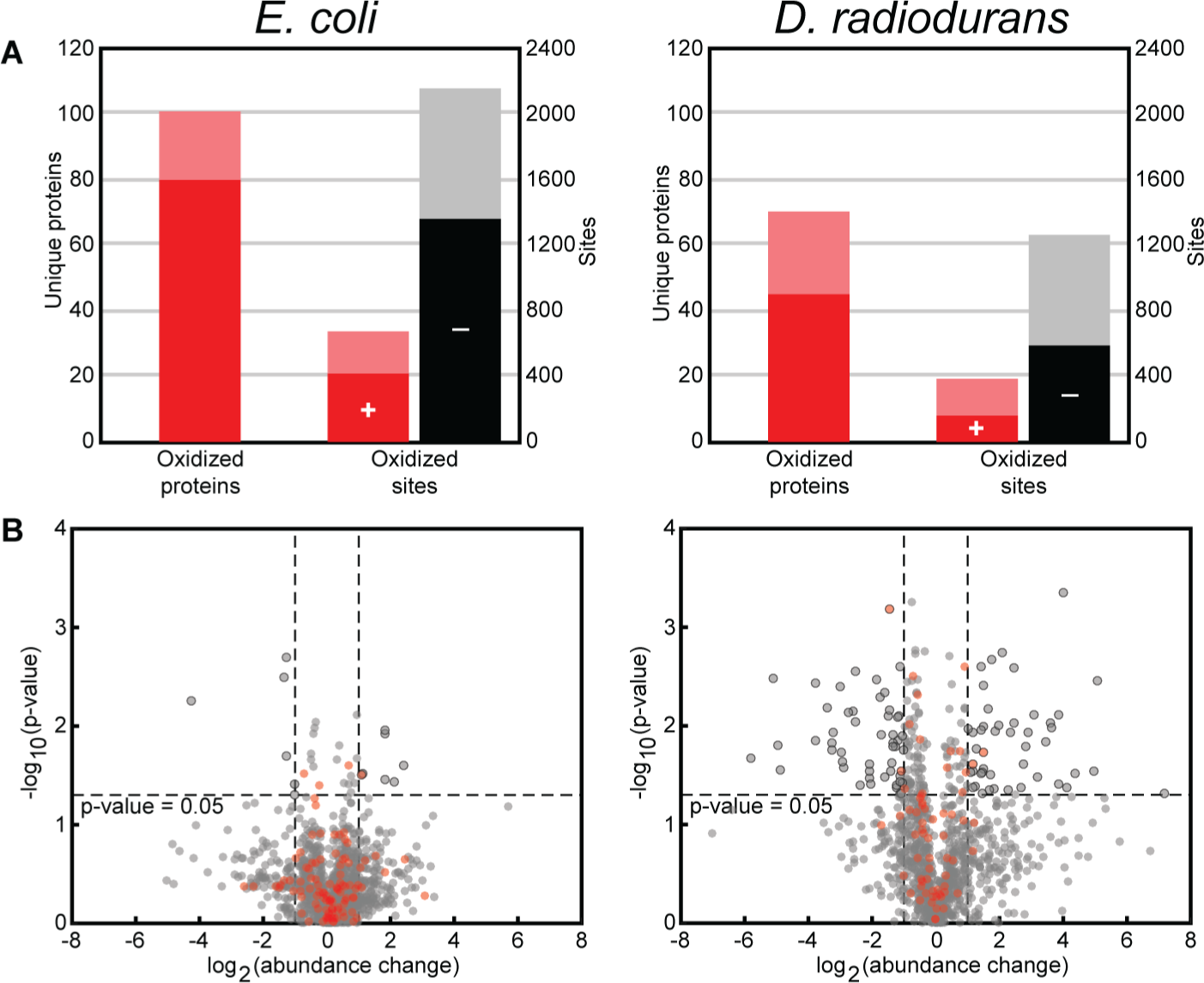
Summary of shotgun redox proteomic data. (A) Carbonyl-bearing proteins detected by shotgun redox proteomic measurement in *E. coli* and *D. radiodurans*. The left axis is the number of sequence-unique proteins detected as oxidized. The right axis is the number of sites in total detected as oxidized (+) or not oxidized (-) in peptides bearing at least one carbonyl. Dark red = oxidized proteins and sites detected in unirradiated samples. Light red = oxidized proteins and sites detected only in irradiated samples. Black = non-oxidized RKPT residues detected in oxidized peptides in unirradiated samples. Gray = non-oxidized RKPT residues detected in oxidized peptides only in irradiated samples. See also Fig. S1. (B) Volcano plots for relative protein abundance changes in *E. coli* (left) and *D. radiodurans* (right) after irradiation. Black-circled points are those proteins with significant changes (t-test p-value < 0.05) of >2-fold or <0.5-fold. Red points are proteins with at least one oxidized peptide detected. Fold change and p-value cutoffs considered for significance are indicated by dashed lines. See also Fig. S2.

Importantly, this resulted in differential relative protein abundances due specifically to oxidative damage (Materials and Methods), distinguishing our results from previous proteomic studies.

As expected (*2*), we observed carbonylation of more proteins in *E. coli* (∼700 CS in 101 of 1206 identified proteins) than in *D. radiodurans* (∼400 CS in 65 of 1125 identified proteins) (Fig. 2A and Table S1). *D. radiodurans* showed similar detection rates to that in *Photobacterium angustum* exposed to UVB (62 carbonylated proteins of 1221 identified) using the same redox proteomic technique (*16*). The lesser total protein carbonylation in *D. radiodurans* was likely due to its effective ROS detoxification mechanisms (*24*). CS saturation curves suggest the fewer detected carbonylation events in *D. radiodurans* account for a greater percent coverage of all *in vivo* events than is the case for *E. coli* (85% and 27%, respectively) (Fig. S2B), in agreement with the difference in oxidation sensitivity between these species.

**Table 1.**
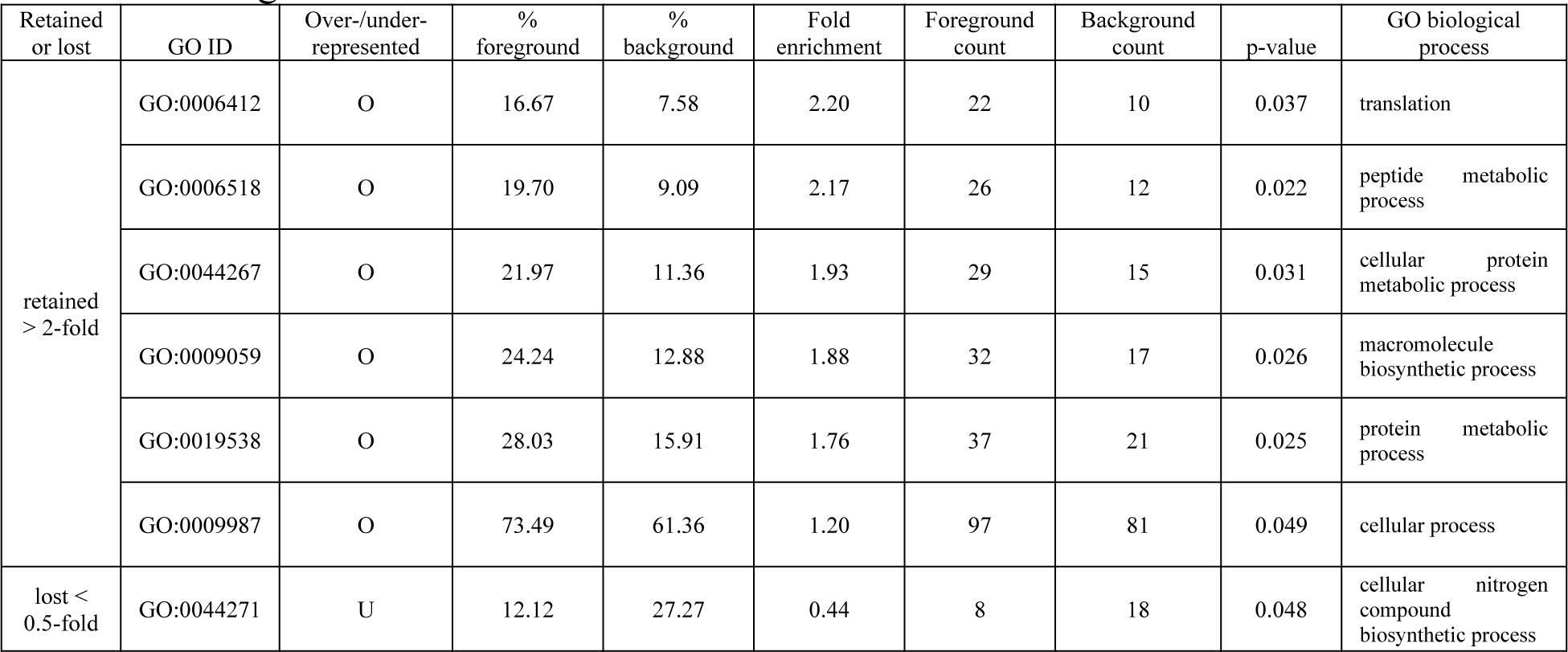
Gene Ontology terms enriched among *D. radiodurans* proteins with high relative abundance change.

Relative quantification provided clear evidence of contrasting differential protein damage distinguishing these organisms (Fig. 2B and Table S2). Although only 0.5% of the identified proteome of *E. coli* showed significant >2-fold differential relative abundance (t-test p-value < 0.05), 15% of *E. coli* proteins overall showed >2-fold changes albeit with higher variability across replicates. In *D. radiodurans*, 8% of proteins significantly changed in relative abundance by >2-fold; the magnitude of change was greater on average with lower variability than in *E. coli*. Carbonylated proteins decreased in relative abundance more than other proteins in *D. radiodurans* (t-test p-value = 0.0016) but less significantly in *E. coli* (t-test p-value = 0.13).

Hence, although *E. coli* accumulated more overall oxidative damage, its distribution is more stochastic across distinct protein species, providing evidence of protein-specific mechanisms for protection from oxidation in *D. radiodurans* that are absent in *E. coli*.

Broad functional characterization of proteins with substantial relative abundance change (<0.5-fold or >2-fold) was carried out by Gene Ontology (GO) biological process term enrichment analysis with protein abundance correction (*25*). These proteins in *E. coli* exhibited no significantly over- or underrepresented GO annotations. In contrast, *D. radiodurans* proteins with >2-fold relative increase were overrepresented by proteins involved in translation and broader protein metabolism (Table 1), including many ribosomal subunits. Additionally, *D. radiodurans* proteins with <0.5-fold change underrepresented proteins involved in nitrogen compound biosynthesis, indirectly implicating the importance of amino acid and nucleotide synthesis. Therefore, resistance to protein oxidation in *D. radiodurans* preferentially protects the critical process of proteome regeneration under oxidative stress.

### 2. Amino acid composition protects against oxidative damage

Although the relative frequency of carbonylated RKTP residues broadly agreed with previous studies (*16, 26*), we found lysine to be as susceptible as proline to oxidation under γ-irradiation (Fig. 3A) in *D. radiodurans* (ratio 1.77 versus 1.66) and to a lesser extent in *E. coli* (ratio 1.17 versus 1.43). Protein oxidation by natively-generated ROS in eukaryotes (*26*) and UV irradiation in *P. angustum* (*16*) both indicated proline as the most ROS-susceptible of RKPT and lysine as not especially or least susceptible, respectively. Proline carbonylation often leads to polypeptide self-cleavage, which may explain the relatively low proline content of bacterial ribosomal versus non-ribosomal proteins (*27*), an evolutionary adaptation contributing to protection of translation against oxidative stress. In contrast, lysine, found incorporated into proteins much more frequently, lacks a similar mechanism for self-cleavage upon oxidation. The more complex role of lysine in oxidative stress is discussed below. Selective amino acid composition is a major adaptation organisms have evolved to thrive in diverse environmental niches (*28*). Comparing compositions between expressed proteomes of *coli* and *D. radiodurans* under permissive conditions (Fig. 3B) revealed significant differences among oxidizable amino acids. Lysine and arginine, both positively charged at physiological pH, differ in ROS-susceptibility and exhibited significant usage differences. While highly-susceptible lysine was found to be less frequently used in *D. radiodurans*, less susceptible arginine was overrepresented instead (0.71-fold and 1.57-fold, respectively). Reversibly-oxidizable sulfur-containing amino acids, cysteine and methionine, were rare in both species, but significantly less prevalent in *D. radiodurans* under permissive conditions (0.53-fold and 0.17-fold, respectively). Surface methionines and cysteines help protect proteins from oxidative damage in many organisms due to their own reversible oxidation (*29*). However, cysteine and methionine are metabolically expensive (i.e. stoichiometrically consume the most ATP) for bacterial synthesis (*30*), and *D. radiodurans* is auxotrophic for methionine (*31*), which may explain their significantly lower prevalence in slower-growing *D. radiodurans* despite expected benefits for resistance. Tryptophan and tyrosine, two metabolically inexpensive amino acids that function as integrated antioxidants in some proteins (*32*), were significantly more abundant in *D. radiodurans* than in *E. coli* (both ∼1.3-fold).

**Fig. 3.**
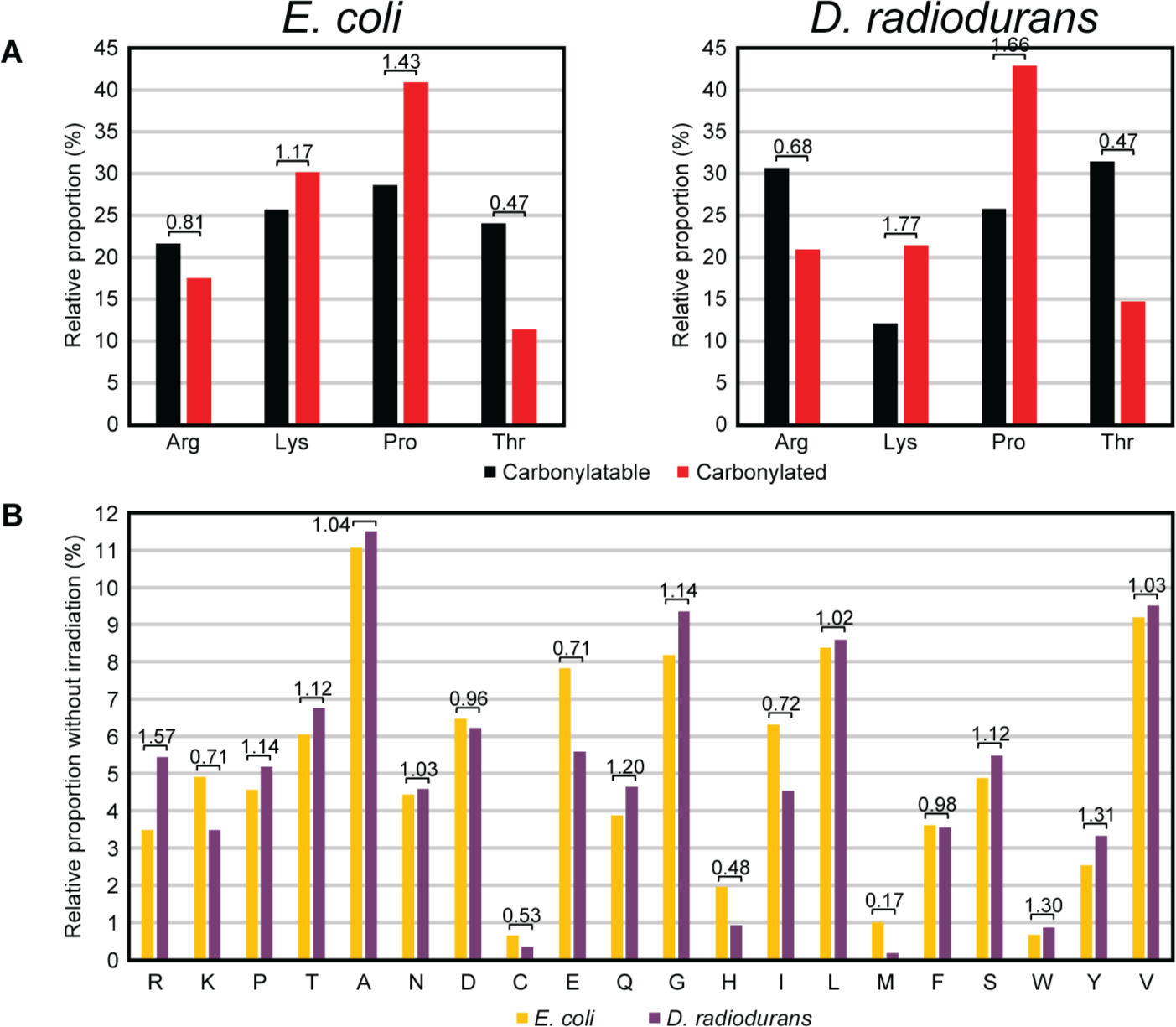
Amino acid prevalence before and after oxidation. (A) Prevalence of individual RKPT residues and prevalence of oxidized form in experimentally measured peptides. Ratios are given above each pair of bars. All proportions are significantly different between each RKPT and their respective oxidation state by z-test (p-values < 0.01), meaning oxidized proportions are not determined simply by relative prevalence of RKPT. See also Fig. S1. (B) Prevalence of all canonical amino acids before irradiation of *E. coli* and *D. radiodurans*. Ratios are given above each pair of bars. All proportions are significantly different between species by z-test (p-values < 0.01). See also Fig. S2 and S3.

To evaluate the impact of oxidative stress on amino acid prevalence in identified proteins, we compared changes in amino acid composition after γ-irradiation of *E. coli* and *D. radiodurans* (Fig. S3). While only 7 amino acids significantly changed in *E. coli*, 16 significantly changed in *D. radiodurans* and to a greater magnitude. The greatest decrease among RKPT was lysine in both species, further supporting that incorporated lysine is an important mediator of protein oxidative damage under γ-irradiation. Lysine can sometimes be exchanged for histidine in proteins and still preserve protein function as shown in synthetic mutational studies (*33*).

Notably, relative histidine prevalence increased modestly (+2%) in *E. coli* and significantly (+11%) in *D. radiodurans* after irradiation, suggesting that *D. radiodurans* has evolved proteins that are more composed of non-carbonylatable histidine rather than lysine as another protein-intrinsic protection mechanism. Indeed, across sequences of functional orthologs and isozymes in these species (Fig. S4) we found 10% greater histidine composition in *D. radiodurans* than in *E. coli* as a fraction of total histidine and lysine (paired t-test p-value < 6×10^-60^). Following irradiation, tyrosine prevalence significantly increased in *E. coli* (+4%) and in *D. radiodurans* (+8%), and cysteine increased significantly (+18%) only in *D. radiodurans*. The most significant decrease in *E. coli* (-13%) and increase in *D. radiodurans* (+45%) was for methionine. This contrast suggests a more efficient methionine sulfoxide reductase system under oxidative stress in *D. radiodurans*. All together, these results establish that protein-intrinsic properties, even in primary structure, differ between *E. coli* and *D. radiodurans* and affect which proteins withstand the onslaught of ROS-induced oxidative damage.

### 3. Structure- and sequence-based model predicts protein vulnerability to oxidation

#### 3.1. Structure-based molecular feature engineering

The computational phase of this study (Fig. 1B) involved proteome-wide derivation of 3D structures to investigate molecular properties contributing to ROS susceptibility (Fig. 4A and Materials and Methods). Due to incomplete proteome coverage by crystal structures (<3% for *D. radiodurans* proteins), computation of molecular features required high-throughput modeling of single-chain proteins, which we performed *de novo* for *D. radiodurans* (Table S3) and used published models for *E. coli* (*23, 34*). The challenge of deriving *D. radiodurans* proteins by available modeling strategies is summarized in Fig. S5A. The best-representative model from alternative methods (Table S4) for each protein was selected using multiple structure quality metrics (Table S5). Models generally evaluated comparably to crystal structures for *D. radiodurans* proteins by these metrics (Fig. S5B). Best-representative models were obtained for >95% of *D. radiodurans* proteins (Fig. S5C), most commonly resulting from I-TASSER (*23*) or ProtMod (http://protmod.godziklab.org/protmod-cgi/protModHome.pl). Future replacement with higher-quality models or experimentally-determined structures could improve the performance of our algorithm.

**Fig. 4.**
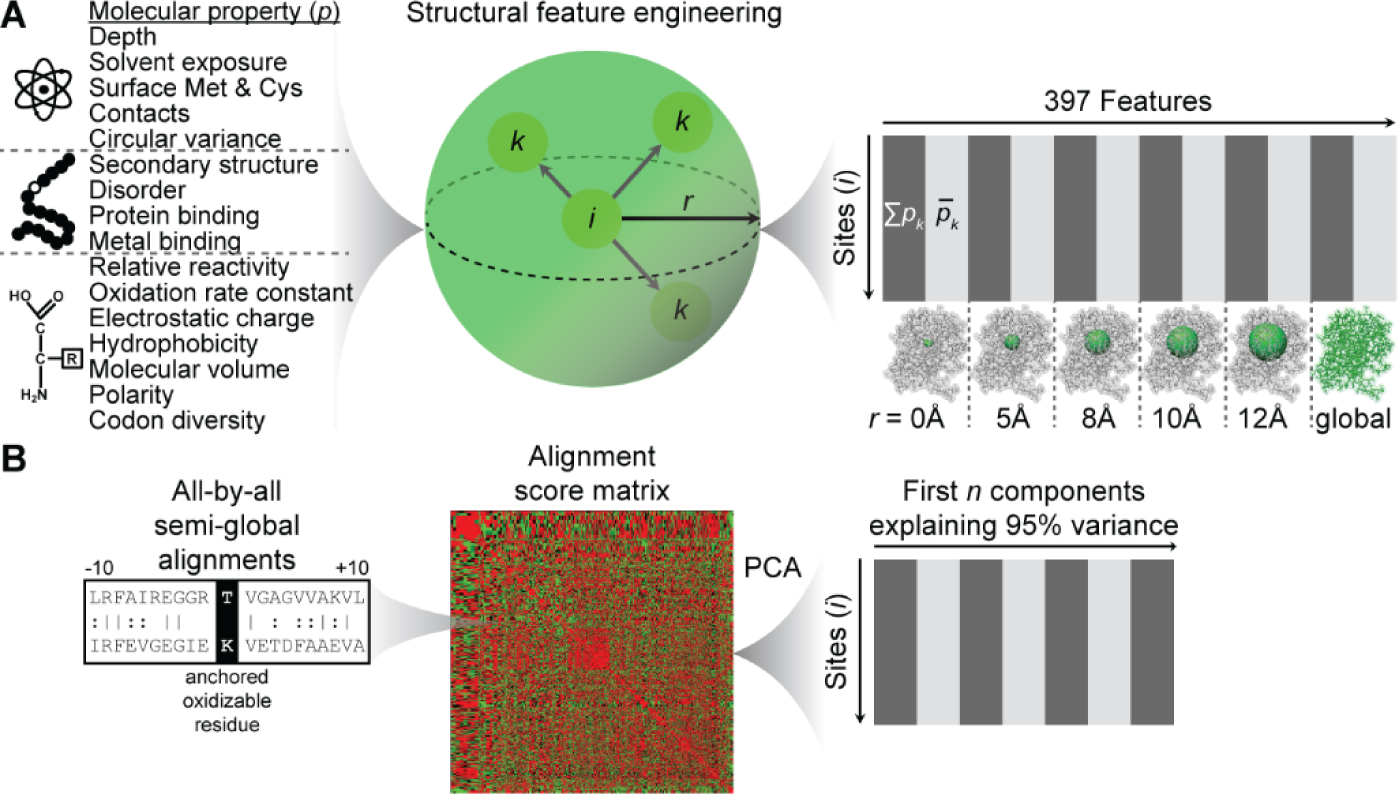
Feature engineering. (A) Three-dimensional feature engineering from molecular properties. Initial properties that can be determined only with an atomic resolution structure, in the context of an amino acid sequence, or that depend only on amino acid identity are denoted at left. This property list is a non-redundant abbreviated set of all properties considered (see Table S6 and Materials and Methods for full detail). Columns of the feature matrix at right are alternating property sums and means at spatial scales denoted below matrix. *p* = a molecular property; *i* = RKPT residue; *k* = neighbor residues of *i*; *r* = radius length. See also Fig. S5. (B) Sequence-homology-based features for machine learning were derived by performing sequence alignments of all RKPT sites (+/- 10 residues) anchored at the central residue to compute alignment scores that were then reduced to a computationally manageable number of features by principal component analysis (PCA).

We engineered for the first time molecular features at multiple spatial scales using 3D structures (Fig. 4A, Table S6, and Materials and Methods) to predict carbonylation. Features were computed with respect to all RKPT across *D. radiodurans* and *E. coli* proteomes. These features quantitatively summarize the molecular environment of oxidizable sites. Statistical summaries of local structural properties were computed as the sums and means of canonical property values for neighboring residues within multiple radii to account for a gradient of scales. This feature engineering strategy enabled incorporation of more molecular properties and with spatial dimensionality than possible using sequences alone to represent proteins.

#### 3.2. Combining structure- and sequence-based approaches for machine learning

In addition to structure-derived features, we implemented simple sequence-alignment-based feature engineering to predict oxidation sites (Fig. 4B). We defined a local neighborhood centered on each RKPT covered by oxidized peptides in our proteomic data and performed all-by-all pairwise sequence alignments of these regions, using the alignment score matrix as potential predictive features. This alignment-based approach is agnostic to specific sequence motifs while still leveraging any useful local sequence homology across CS.

All RKPT from oxidized peptides (Table S1) were mapped to respective protein structure and sequence to assign carbonylated and non-carbonylated residues (Table S7). Unlike previous CS prediction efforts (*17–19*), we did not assume that any given RKPT is deterministically oxidized or not. Protein oxidation is an inherently stochastic process. Therefore, we took a probabilistic approach and used all of the oxidized peptide data regardless of site redundancy or occurrence as carbonylated in one peptide but non-carbonylated in another. Previous approaches also often sampled unmodified RKPT across all detected peptides, oxidized or not, to define negatives for training. Compared to non-oxidized peptides, unmodified RKPT on peptides bearing a carbonyl on another residue better-represent negative data because it is certain that those molecules were directly exposed to ROS yet did not react with ROS.

Independent probability estimators for CS were trained by logistic regression using structure-based features and sequence-based features and then combined into a stacked model. Each independent model and the stacked model were evaluated by leave-1-out validation and their performance quantified by receiver operating characteristic (ROC) analysis (Fig. 5A). At the residue scale, our stacked model outperformed (AUCnorm = 0.73) each of its structure- and sequence-based components. Shuffling each feature before training yielded random performance (AUCnorm = 0.54), strongly supporting the predictive power of our engineered features. We also evaluated performance of our model for predicting protein-scale vulnerability to oxidation (Fig. 5B) by calculating an oxidation site enrichment metric. Predicted oxidation enrichments for training set proteins strongly rank correlate with oxidation enrichments derived from measured oxidized peptides (ρ = 0.82, p-value = 1.3×10^-22^ for *E. coli* and ρ = 0.87, p-value = 7.2×10^-21^ for *D. radiodurans*), signifying that our model can predict relative propensity to carbonylation of different protein species. Due to prioritized sensitivity, our model tends to predict higher oxidation enrichment values than derived experimentally (1.9-fold on average for *E. coli* and 1.7-fold for *D. radiodurans*), but these predicted enrichment values are plausible given the fact that *in vivo* oxidation events are under-sampled experimentally (Fig. S2B).

**Fig. 5.**
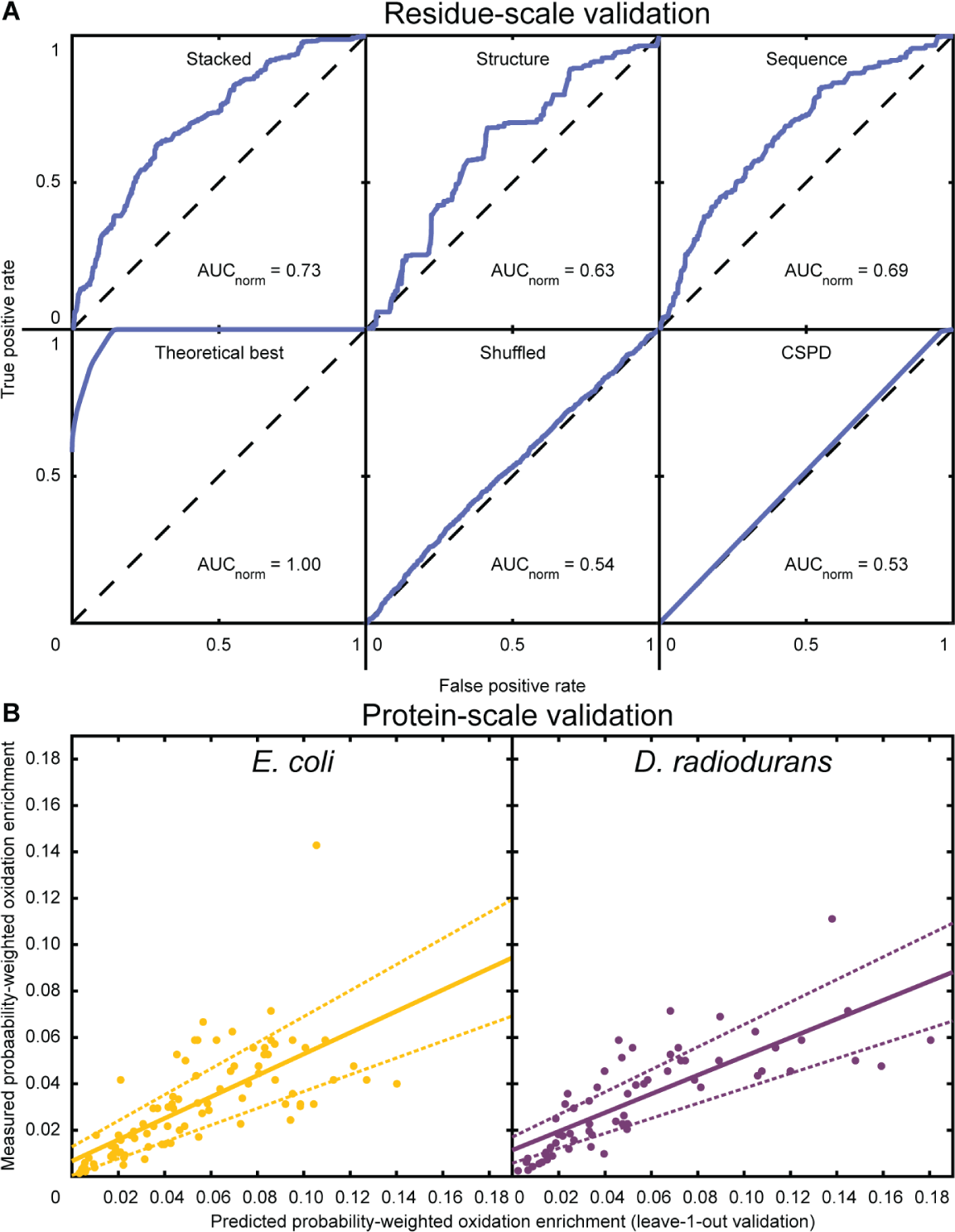
Multi-scale validation of protein oxidation predictor. (A) Residue-scale validation: Receiver operating characteristic (ROC) curves for oxidation site predictors derived by leave-1-out validation. The dashed black line at y=x corresponds to performance expected by chance. Top left = final predictor trained by stacking structure- and sequence-based models. Top middle = predictor trained only on structure-based features. Top right = predictor trained only on sequence-based features. Bottom left = theoretical maximum predictive power for a probability estimator (AUC = 0.98). Bottom middle = same algorithm as used for final predictor but with all features shuffled beforehand. Bottom right = CSPD model developed using metal-catalyzed oxidation (MCO) site data from *E. coli*. See also Fig. S5 and S6. (B) Protein-scale validation: Comparison between predicted oxidation site enrichment from leave-1-out validation to oxidation site enrichment computed from all oxidized peptides measured for *E. coli* (left) and *D. radiodurans* (right). Each point represents a different protein species. Predicted probability-weighted oxidation site enrichment = (sum of oxidation probabilities across training set sites)/number of residues in corresponding peptides from experiments. Experimentally measured probability-weighted oxidation site enrichment = (sum of empirical oxidation probabilities across training set sites)/number of residues in corresponding peptides from experiments. The solid line is the fitted regression line, and dashed lines indicate the boundaries of the 95% confidence interval.

#### 3.3. Molecular properties explain vulnerability to oxidation

Although we included ∼400 structure-based features in the modeling, only 7 of the logistic regression coefficients were non-zero: relative reactivity with ROS (reactivity_res), codon diversity, whether the RKPT site was a threonine residue, molecular volume, local solvent accessible surface area, local positive charge, and local lysine residues. Codon diversity (AAindCodonDiv_res) itself is unlikely to be causal. Instead, this feature has the same rank order as oxidation prevalence in *D. radiodurans* from our experiments (Fig. 3A) and is therefore a fortuitous proxy for γ-specific reactivity. Threonine is by far the least frequently carbonylated of RKPT in both species (Fig. 3A), and inclusion of this feature (Thr_res) in our model reflects this lower propensity to reaction with ROS.

Aside from the reactivity features differentiating RKPT, all other explanatory properties for ROS susceptibility derived from 3D structures (Fig. 6). Accessibility to ROS promotes oxidation (Fig. 6A). The lower the molecular volume of a residue (AAindMolVol_res), the more likely it is oxidized due to lower steric effects. Similarly, lower local surface area (areaSAS_5A_sum) surrounding a near-surface site indicates less likelihood of shielding by surrounding structure, such as the protrusion in Fig. 6D. Local positive charges (posCharge_8A_sum) promote oxidation by attracting negatively charged superoxide radicals (Fig. 6B). Co-localization of highly reactive sites may cause progressive protein misfolding, exposing neighboring residues to ROS (*17*) (Fig. 6C). In our model, neighboring lysine residues (Lys_8A_sum) contribute to the probability of oxidation, lysine being the most prevalently oxidized RKPT under γ-irradiation in our data (Fig. 3A). Polarity leading to solubility of lysine-rich regions could also contribute to this effect. Sites without neighboring lysines are less likely to be oxidized (Fig. 6D).

**Fig. 6.**
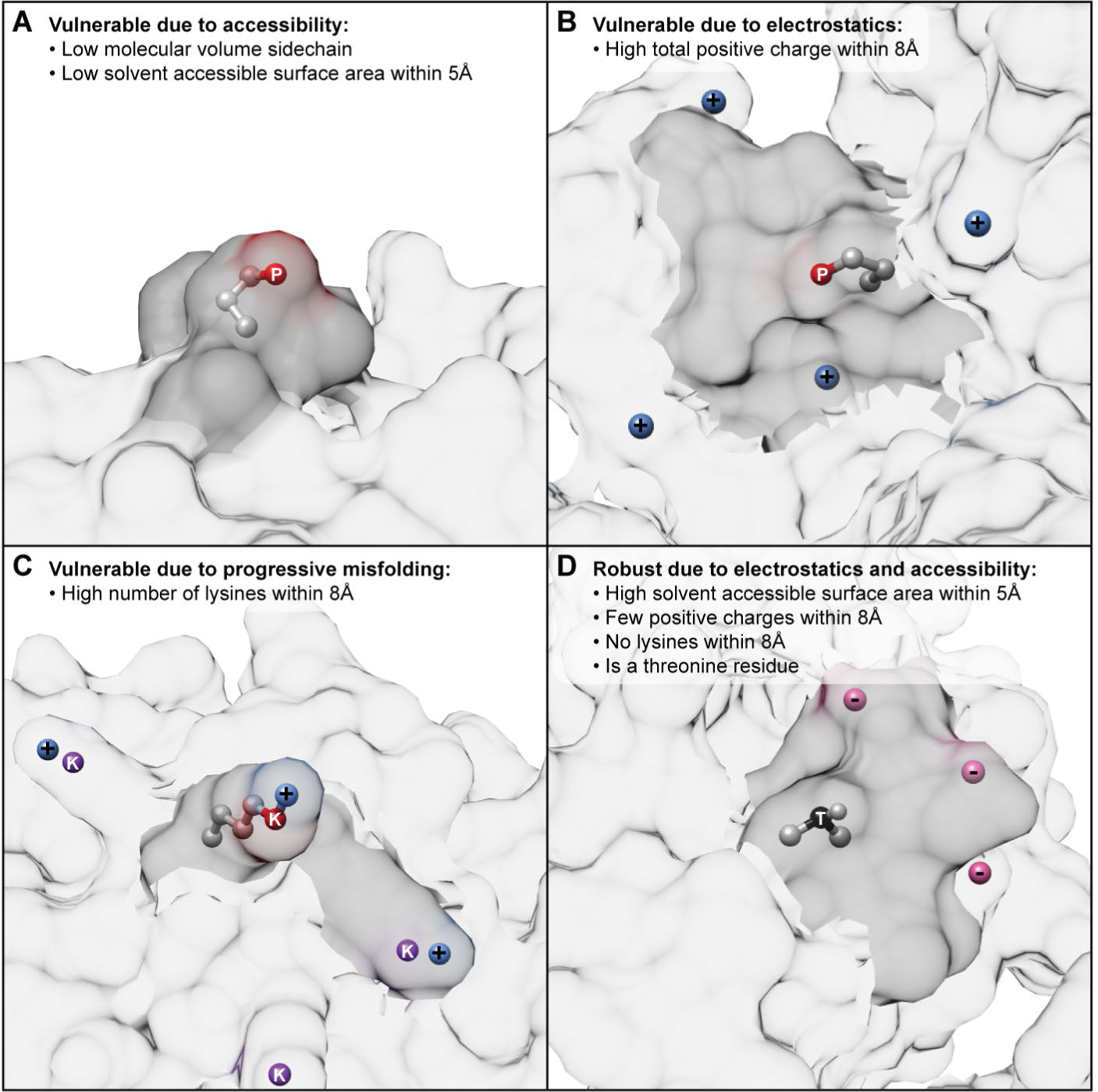
Molecular properties predicting protein vulnerability to oxidation. Example sites prone to oxidation (A) DRA0302_P252, (B) DR0099_P51, and (C) b0911_K411; and example robust site (D)) b3313_P69. All atoms of central RKPT side chains are shown, with oxidizable atomic site in red (predicted and measured oxidized) or black (predicted and measured not oxidized) and labeled with the 1-letter code of the containing amino acid. Positive (blue) and negative (pink) charges within 8Å are labeled. Oxidizable lysine sites (purple) within 8Å are labeled. Molecular surfaces within 5Å of the central oxidizable site are dark gray. See also Fig. S5.

#### 3.4. Our algorithm also extends to prediction of metal-catalyzed oxidation

We applied Carbonylated Site and Protein Detection (CSPD) (*17*) to predict CS across our training set (Fig. 5A). CSPD performance on our data was essentially random (AUCnorm = 0.53). It is important to note that CSPD was developed using metal-catalyzed oxidation (MCO) data from a small set of *E. coli* proteins, and the inability of CSPD to generalize to oxidation from γ-irradiation may be due in part to a difference in effects of each specific source of ROS. Therefore, to more directly compare algorithmic performance we also used our algorithm to train a model predicting MCO using the same redox proteomic data used to develop CSPD (Fig. S6). CSPD showed modest positive performance on this dataset (AUCnorm = 0.58), the discrepancy in previously reported performance owing to our inclusion of all oxidized peptides with carbonylated and non-carbonylated residues defined as described above. We conclude that CSPD was overfitted to the MCO data and depends on the assumption of deterministic protein oxidation and on less strict standards for defining non-carbonylated residues in proteomic data.

Furthermore, our stacked model for MCO prediction performed better (AUCnorm = 0.75) than our γ-induced oxidation model with better synergy in stacking the structure-(AUCnorm = 0.72) and sequence-based (AUCnorm = 0.67) models. This performance difference was likely due to the relatively less diverse products of MCO than γ-induced oxidation. ROS production in MCO is more localized because it depends on the presence of Fe or Cu cations to drive the Fenton reaction and therefore affects a smaller number of proteins than γ-induced oxidation.

Indeed, data from γ-irradiation experiments includes not only CS caused by ROS from water radiolysis but also basal cellular oxidation due to native ROS sources, including MCO and cellular respiration. Thus, CS from γ-irradiation is more diverse and complex than MCO products and more challenging for learning structure and sequence signatures.

### 4. Intra- and interspecies differences in protein oxidation vulnerability

#### 4.1. *D. radiodurans* proteome maintenance is protected from oxidation

Orthologs and isozymes mapped between *E. coli* and *D. radiodurans* (Fig. S4) were compared by their unweighted oxidation enrichment (Fig. 7) as computed from proteome-wide CS prediction in *E. coli* (Table S8) and *D. radiodurans* (Table S9) to reveal functional classes and individual proteins differing in susceptibility between and within these proteomes.

**Fig. 7.**
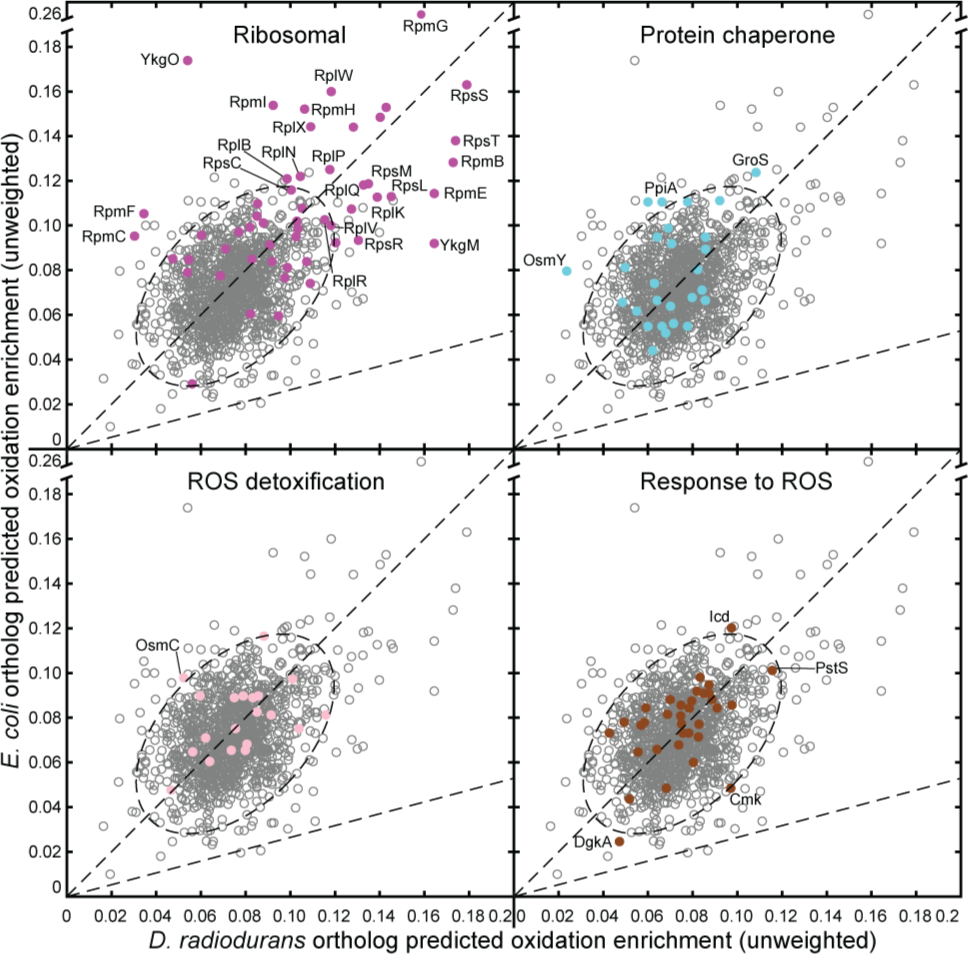
Interspecies comparison of predicted protein vulnerability to oxidation. Each circle represents a distinct protein pair (ortholog or isozyme) between species. Each plot shows identical point values but highlights a different functional class of proteins with relevance to oxidative stress. The y=x diagonal line is a reference to compare intrinsic vulnerability to oxidation between orthologs. The y=x/3.78 diagonal line is a reference to compare combined intrinsic and extrinsic oxidation properties between orthologs. Elliptical dotted line encircles the points falling within 3 standard deviations of the mean coordinates and 3 standard deviations of the distance from reference line y=x, encompassing ∼91% of all data points. This reference region distinguishes outlier points that are distant from the main population. Outliers with associated experimental evidence related to hypersensitivity to oxidative stress are labeled with their protein names. See also Fig. S4, Fig. S7, and Table S10.

Functional classes known to be involved in resistance and recovery from oxidative stress include: ribosomal, ribosomal assembly, translation, protein chaperone, protease and peptidase, amino acid and peptide transport, DNA repair, DNA damage response and regulation of repair, native ROS production, ROS detoxification, ROS response, metal transport, terpenoid synthesis, and polyamine accumulation (Table S10).

Pairwise orthologs were compared based on protein-intrinsic and extrinsic factors contributing to their propensity to carbonylation (Fig. 7). Perpendicular distance to the y=x diagonal represents the relative degree to which one ortholog is intrinsically more or less sensitive given the same ROS dosage on the basis of oxidation enrichment alone. Protein-extrinsic factors, such as the Mn-dependent scavenging system in *D. radiodurans* (*35*) and the antioxidant carotenoid deinoxanthin (*36*), also contribute to interspecies differences in protein oxidation. Such protein-extrinsic factors act broadly by reducing the effective cellular dosage of ROS. An acute gamma dosage of 7kGy, approximately the same as in this study, yielded about 3.78-fold more protein carbonyls in *E. coli* lysate than in *D. radiodurans* (Materials and Methods) due to small molecules removable by dialysis (*2*). Assuming such factors act globally without favoring protection of specific proteins, the degree to which these extrinsic factors differentiate vulnerability to oxidation between orthologs can be modeled in combination with protein-intrinsic factors simply by computing perpendicular distance to the y=x/3.78 diagonal (Fig. 7). By this model, especially susceptible proteins benefit more from an effectively lower dosage of ROS in *D. radiodurans*.

Relative vulnerability to ROS differed between *E. coli* and *D. radiodurans* within particular functional classes (Fig. 7 and Table S10). We predicted the intrinsic susceptibility of *E. coli* ribosomal proteins to be more than 2.4-fold greater than across all-orthologs (p-value = 0.01). Accounting for extrinsic ROS protection predicted ribosomal proteins to be the most favored functional class in *D. radiodurans* over *E. coli* (1.5-fold, p-value = 1.2×10^-26^), in agreement with *D. radiodurans* ribosomal proteins being enriched among those with relative abundance increases after irradiation. Protein chaperones in *E. coli* were predicted on average 1.13-fold more intrinsically vulnerable than in *D. radiodurans* (p-value = 0.02), a difference further distinguished due to being more than 4.5-fold greater than the difference across all-orthologs (p-value = 0.003) and 1.14-fold greater when accounting for extrinsic protection as well (p-value = 0.02). *E. coli* proteins involved in polyamine synthesis and uptake are predicted to be more than 3.7-fold intrinsically vulnerable than across all-orthologs (p-value = 0.04). Revisiting the observation that methionine usage featured prominently in *D. radiodurans* proteins retained after irradiation, we predicted that methionine sulfoxide reductases acting on protein-incorporated methionine MsrB and MsrP are both 1.4-fold more intrinsically sensitive to carbonylation in *E. coli*. MsrP was also in the 94^th^ percentile of proteins benefiting from extrinsic protection in *D. radiodurans*.

#### 4.2. Comparison of interspecies outliers reveals proteins involved in oxidative stress resistance

Many proteins involved in coping with oxidative stress were significant outliers in predicted intrinsic vulnerability to oxidation (Fig. 7 and Table S10). There were 111 orthologous pairs greater than 3 standard deviations of distance from the mean of the distribution or greater than 3 standard deviations away from the mean perpendicular distance from the y=x diagonal.

We grouped these outliers according to three properties: 1) intrinsic sensitivity or robustness compared to the rest of the proteome, 2) comparative intrinsic vulnerability between *D. radiodurans* and *E. coli*, and 3) relative effect of ROS detoxification in *D. radiodurans* over *E. coli* (Fig. S7 and Table S10).

Proteins predicted as significantly more intrinsically or extrinsically protected from ROS in *D. radiodurans* relative to *E. coli* fall into three groups based on the three properties described above. Group 1 proteins were predicted highly oxidation-prone but more protected intrinsically and extrinsically in *D. radiodurans* than in *E. coli*. On average, these 12 proteins were 1.4-fold more oxidation-site enriched in *E. coli* and above the 99^th^ percentile of extrinsic protection in *D. radiodurans*. Of 10 proteins detected in both organisms by proteomics, 8 had more negative γ-induced relative abundance changes in *E. coli* than *D. radiodurans*, with a median *E. coli*-to-*D. radiodurans* ratio of 0.47. Ribosomal subunits comprised 11 of these proteins, 8 of which are essential in *E. coli*. *E. coli* knockouts of *rpmI* (*37*) are hypersensitive to oxidative stress. Overexpression of *rpmG* increases resistance to oxidative stress from mitomycin C (*38*), and GroS overexpression decreases protein carbonyl accumulation (*39*). Seven of these proteins exhibit oxidative-stress-induced expression in *D. radiodurans* (*24, 40*). Group 2 proteins were predicted as similarly intrinsically oxidation-prone in both species but significantly extrinsically protected in *D. radiodurans*. On average, these 22 proteins are above the 86^th^ percentile of extrinsic protection in *D. radiodurans*. Of 13 proteins detected in both organisms by proteomics, 11 showed substantially more positive γ-induced relative abundance changes in *D. radiodurans*. In this group, 13 proteins are ribosomal subunits. In *E. coli*, *pstS* knockouts are hypersensitive to oxidative stress (*41*), *rpsL* mutants have been shown to affect oxidative stress tolerance (*42, 43*), and 13 others are essential genes (*44, 45*). In *D. radiodurans rpsS* and *hupA* knockouts are hypersensitive to oxidative stress (*46*), and overexpression of *rpsS*, *rpsT*, *rplQ*, *rpsM*, *rpmB*, *rplK*, *rpsL*, *thpR*, *rpmE*, *nrdH*, *rplR*, *rplV*, and *rpsR* occurs during oxidative stress (*24, 40*).

Group 3 proteins were predicted significantly more susceptible to oxidation in *E. coli* than in *D. radiodurans*. On average, these 27 proteins were 1.9-fold more oxidation-site enriched in *E. coli* and above the 95^th^ percentile of extrinsic protection in *D. radiodurans*. In *E. coli*, *rpmF* (*37, 41*) and *icd* (*47*) knockouts are hypersensitive to oxidative stress, and *osmY* (*48*) is also involved in oxidative stress resistance. In *D. radiodurans xseB* knockouts are hypersensitive to oxidative stress (*46*), and *adk*, *icd*, *malE*, *osmC*, *ppiA*, *rplB*, *rpmC*, *rpsC*, and *yceI* are highly expressed under oxidative stress (*24, 40, 49*). Higher resistance to oxidation of proteins from these groups sets *D. radiodurans* apart from *E. coli* and delineates transgenes that could serve to increase stress tolerance in *E. coli*.

Interspecies outliers not predicted as significantly more protected from ROS in *D. radiodurans* fall into two groups. Group 4 proteins were predicted as highly intrinsically robust to oxidation in both species and therefore not to benefit substantially from extrinsic protection in *D. radiodurans*. Of these 5 proteins, 3 were more intrinsically vulnerable in *E. coli*, including *secE*, which is essential in *E. coli* (*44*), and *fdx*, which is highly expressed under oxidative stress in *D. radiodurans* (*40*). Group 5 proteins were predicted as significantly more intrinsically vulnerable to oxidation in *D. radiodurans* than in *E. coli*. These 14 functionally diverse proteins include 3 known oxidative-stress-hypersensitive knockout mutants in *D. radiodurans* (*46*); however, all but 2 still lie above y=x/3.78 in Fig. 7, suggesting that extrinsic protection could still compensate for intrinsic vulnerability differences between these species.

## Conclusion

In this study, we successfully developed a highly integrative systems biology approach that predicts protein targets of oxidative stress and offers mechanistic explanations for cellular phenotypes at multiple biological scales spanning amino acid residues, protein molecules, and protein functional classes. Multiple lines of evidence support that the susceptibility of ribosomal proteins to ROS is strongly differentiated from the rest of the proteome and plays a key role in the radioresistance of *D. radiodurans*. We have provided substantial evidence that intrinsic properties of proteins affect vulnerability to oxidative stress. Our model suggests that a combination of protein-intrinsic properties and global ROS-detoxification implicates vital proteins for resisting oxidative stress in *D. radiodurans* and explains targeted patterns in relative abundance changes following irradiation that are not observable in *E. coli*. Those *D. radiodurans* proteins predicted to rely primarily on intrinsic properties to avoid oxidation are candidate transgenes to confer resistance to more sensitive species. Amino acid usage differentiating *D. radiodurans* from *E. coli* and molecular properties predictive of protein carbonylation comprise a set of design principles that may be used to control ROS-tolerance in synthetic protein engineering efforts. The analytical strategy that we have established could be applied not only to the study of oxidative stress in other systems (e.g. human disease, aging, and manned space exploration) but also to other forms of post-translational modification and more broadly to molecular properties of proteins, extending and enriching proteomic analysis.

## Acknowledgments

Roger Chang especially thanks Dr. Pamela Silver (Harvard) who as his Postdoctoral Advisor allowed and supported this unique, independently conceived effort. As such, Dr. Silver provided laboratory resources, feedback on the work, comments on the manuscript and full support of this independent publication. We also thank Dr. Debora Marks (Harvard) for advice on machine learning, data analysis and feedback on the manuscript; Dr. James Collins (MIT) for facilitating γ-irradiator access; Dr. James Weaver (Wyss Institute) for fabricating the irradiator rack; and Dr. Fatma Elzahraa Eid (Broad Institute) for proofreading the manuscript. Computational modeling was enabled by the HMS Orchestra high performance compute cluster.

## Author contributions

R.L.C. and S.M.-S. conceived, supervised, developed experimental and analytical methods for, performed experiments, and acquired funding for this study; R.L.C., M.C.R., J.W.S., and A.R.O. developed software and performed computations; R.L.C., M.C.R., and S.M.-S. performed statistical analysis and validations; Z.L., Y.A.C., R.W., A.G., and S.M.-S. provided study materials and computing resources; R.L.C., M.C.R., J.W.S., Z.L., A.R.O., and S.M.-S. performed data curation; R.L.C., M.C.R., J.W.S., Z.L., Y.A.C., A.R.O., and S.M.-S. wrote the manuscript

## Data and materials availability

Raw proteomic mass spectrometry data for this study is deposited in the Mendeley database (http://dx.doi.org/10.17632/jx52c2xgfk.1). Code for feature computation from structures is available in GitHub (https://github.com/julianstanley/ProteinFeatures.git). All other data is available in the main text or the supplementary materials.

## Materials and Methods

### Redox proteomics

#### Reagent preparation

TGY medium (1 L)

- 5 g tryptone
- 5 g yeast extract
- 1 g glucose
- 1 g potassium phosphate dibasic (K2HPO4)
- (20 g agar for solid medium)
- Fill to 1 L with milliQ water.
- Mixture pH should be 7.0 without, but if needed, adjust to pH 7.0 with 1 N NaOH.
- Sterilize by autoclaving.

LB medium (1 L)

- 10 g tryptone
- 5 g yeast extract
- 10 g NaCl
- (20 g agar for solid medium)
- Fill to 1 L with milliQ water.
- Adjust to pH 7.0 with 1 N NaOH.
- Sterilize by autoclaving.

Harvest buffer (for ∼40 samples)

- 1.2037 g anhydrous MgSO4 (10 mM working concentration)
- Fill to 1 L with milliQ water.
- Mix by swirling until dissolved, and store at 4°C.

Lysis buffer (for ∼130 samples)

- 95.32 mg 4-(2-hydroxyethyl)-1-piperazineethanesulfonic acid (HEPES) (10 mM working concentration)
- 14.41 g urea (6 M working concentration)
- 6.090 g thiourea (TU) (2 M working concentration)
- Fill to 40 mL with BioRad ReadyPrep™ Proteomics Grade Water.
- Dissolve with magnetic stirrer, and adjust pH to 8.0.
- Transfer to a 50 mL polypropylene tube.
- Mix well again by vortexing, and store at 4°C.

7X Protease inhibitor cocktail (for ∼30 samples)

- Transfer one 25 mg cOmplete™, Mini, EDTA-free Protease Inhibitor Cocktail tablet to a 2 mL microfuge tube.
- Dissolve tablet in 1.5 mL BioRad ReadyPrep™ Proteomics Grade Water.
- Store at -20°C. Stable for 12 weeks after preparation.

Derivatization buffer (for ∼130 samples)

- 79.3 mg dinitrophenylhydrazine (DNPH) (10 mM working concentration)
- Transfer to a 50 mL polypropylene tube.
- Fill to 40 mL with 2 M HCl.
- Mix well by vortexing, wrap tube in foil to protect from light, and store at 4°C.

Hydrazone stabilization buffer (for ∼130 samples)

- 75.4 mg sodium cyanoborohydride (NaBH3CN) (30 mM working concentration)
- 4 mL 10X PBS
- Fill to 40 mL with BioRad ReadyPrep™ Proteomics Grade Water.
- Dissolve with magnetic stirrer, and adjust pH to 7.4.
- Transfer to a 50 mL polypropylene tube.
- Mix well by vortexing, and store at 4°C.

Protein precipitation buffer (for ∼220 samples)

- 10 mL 100% trichloroacetic acid (TCA)
- Store in a small glass bottle at 4°C.

Wash buffer (for ∼20 samples)

- 20 mL 100% ethanol (EtOH)
- 20 mL 100% ethyl acetate (EtAc)
- Transfer to 50 mL polypropylene tube, mix well by vortexing, and store at 4°C.

Protein resolubilization buffer (for ∼360 samples)

- 22.93 g guanidine hydrochloride (GuHCl) (6 M working concentration).
- 2.7217 g monobasic potassium phosphate (KH2PO4) (0.5 M working concentration)
- Fill to 40 mL with BioRad ReadyPrep™ Proteomics Grade Water.
- Dissolve with magnetic stirrer, and adjust pH to 7.5 with 10 M KOH.
- Transfer to 50 mL polypropylene tube, mix well by vortexing, and store at 4°C.

10X Tris-glycine SDS-PAGE running buffer, pH 8.3 (1 L)

- 30.30 g Tris base (250 mM working concentration)
- 144.10 g glycine (1.92 M working concentration)
- 50 mL 20% SDS (1% working concentration). Warm SDS in 37°C water bath if any undissolved precipitate is visible.
- Fill to 1 L with distilled water, mix well, and store at room temperature.
- Do not adjust pH. Intrinsic pH should be 8.3.
- Add 100 mL 10X running buffer to 900 mL distilled water for use (1X working concentration).

3X SDS-PAGE loading buffer (for ∼4,000 samples)

- 2.4 mL 1 M Tris-HCl (pH 6.8)
- 3 mL 20% SDS. Warm SDS in 37°C water bath if any undissolved precipitate is visible.
- 6 mL 50% glycerol
- 1.6 mL 2-mercaptoethanol (BME)
- 6 mg bromophenol blue
- 7 mL distilled water
- Combine in a 50 mL polypropylene tube, and mix well by vortexing.
- Aliquot into 1 mL volumes and store at 4°C.

Stain solution (300 mL, reusable)

- 1.25 g Coomassie R-250 (i.e. Brilliant Blue R)
- 25 mL 100% methanol
- 50 mL 100% (glacial) acetic acid
- 225 mL distilled water
- Mix well by swirling, and store at room temperature.

Destain solution (1 L, single use)

- 200 mL 100% (glacial) acetic acid
- Fill to 1 L with tap water, and mix well.

Reducing buffer (for ∼160 samples)

- Prepare immediately prior to use.
- Open a sealed 0.5 mL ampule of 1M dithiothreitol (DTT) (0.25 M working concentration).
- Transfer to a 2 mL microfuge tube.
- 1.5 mL BioRad ReadyPrep™ Proteomics Grade Water
- Mix by pipetting, and keep on ice until use.

Alkylation buffer (for 12 samples)

- Prepare immediately prior to use.
- 37.0 mg iodoacetamide (IAA) (0.4 M working concentration)
- Transfer to a 1.5 mL microcentrifuge tube.
- 0.5 mL BioRad ReadyPrep™ Proteomics Grade Water
- Mix by pipetting, wrap tube in foil to protect from light, and keep on ice until use.

Trypsin reaction buffer (11.11 mL, for ∼75 reactions)

- Add 10 mL of BioRad ReadyPrep™ Proteomics Grade Water to a bottle of Trypsin Reaction Buffer (Sigma-Aldrich Catalog Number R3527) (40 mM ammonium bicarbonate working concentration).
- Add 1 mL of Biotech Grade Acetonitrile (Sigma-Aldrich Catalog Number 494445) (9% working concentration.
- Intrinsic pH is 8.2. Store at 4°C. Stable for 1 month after preparation.

#### Gamma-irradiator culture tube rack

In order to irradiate bacterial cultures in the GC-220E ^60^Co γ-irradiator, a custom culture tube rack was designed and fabricated by 3D printing. The design requirements for the rack were that it:

- Hold six 120-mL samples in leak-proof culture tubes, triplicates of two conditions
- Be made of γ-resistant but non-shielding materials
- Hold enough ice and provide insulation to keep samples near 0°C for ≥ 2 h
- Fit inside the GC-220E sample chamber (I.D. = 15.2 cm × H = 20 cm)
- Provide for radially symmetric sample distribution for even dosing

Six 170-mL pyrex culture tubes (O.D. = 38 mm × L = 200 mm) with screw caps (Corning® 9825-38) were shortened so that the height of the loaded rack would remain below 20 cm. A diamond cutter was used to shorten the pyrex tubes at the open end, and a wet orbital sander was used to shorten the screw caps at the open end and smooth the cut end of the pyrex tubes to fit the rack. The pyrex and resin caps are resistant to repeated acute doses of γ-radiation (http://www.sterigenics.com/services/medical_sterilization/contract_sterilization/material_consid eration irradiation_processing.pdf), although the pyrex slightly discolors upon initial irradiation.

The rack was printed using a compound primarily composed of acrylic and polyacrylate material (Stratasys® Objet RGD515). Acrylic and polyacrylate exhibit high radiation stability, up to 100 kGy with repeated exposures (http://www.nordion.com/wp-content/uploads/2014/10/GT_Gamma_Compatible_Materials.pdf). The culture tubes are held in place by hemispherical recesses in the base. The rack has space to add ice through a capped opening at the top using a funnel, sufficient to keep the samples near 0°C for up to 5 h at room temperature and at least 2 h in the GC-220E with a dose rate of 60 Gy/min. The top of the rack is removable for easy cleaning and has a key slot for the alignment of holes in the lid with hemispherical recesses in the base. STL files containing the design data required for 3D printing are available in Files S1 and S2.

#### Bacterial culture

Day 1:

*D. radiodurans* R1 (ATCC® R™) was recovered in triplicate from frozen glycerol stocks for 24 h in 4 mL TGY liquid medium with 0.2 μg/mL ciprofloxacin to prevent contamination during the slow recovery period. Ciprofloxacin is highly bacteriostatic to *E. coli* and many other bacteria but not *D. radiodurans* at this concentration; because our lab cultures multiple faster-growing species in the same incubator, we took this measure only during the recovery from frozen stock. Without ciprofloxacin, we frequently observed our *D. radiodurans* cultures being overtaken by contaminants at this stage. Recovery cultures were grown aerobically at 30°C in a rotary shaker (130 rpm).

Day 2:

*D. radiodurans* replicates were back-diluted 1:1000 in 250 mL room-temperature fresh TGY (no ciprofloxacin) in 500 mL flasks, and growth was continued aerobically overnight at 30°C. *E. coli* K-12 MG1655 (ATCC® 700926™) was recovered in triplicate from frozen stocks and grown aerobically at 37°C in a rotary shaker (130 rpm) overnight in 4 mL LB.

Day 3:

*E. coli* cultures were back-diluted 1:1000 in 250 mL fresh LB in 500 mL flasks, and growth was continued aerobically at 37°C, monitoring optical density at 600 nm (OD600). Growth of *D. radiodurans* (OD600) was checked every 15 min starting 12 h after back-dilution from the previous evening.

*E. coli* and *D. radiodurans* cultures were harvested at mid-log phase (OD600 = 0.5 to 0.6, 3.5 h after back-dilution for *E. coli*; OD600 = 0.8, 13.25 h after back-dilution for *D. radiodurans*). It should be noted that *E. coli* and *D. radiodurans* have different optimal nutrient media, growth temperatures, cell morphologies, and growth rates. Cultures were harvested by transferring 120 mL volumes into 250 mL pyrex bottles with screw caps (for unirradiated controls) or custom-cut 170 mL pyrex culture tubes with screw caps (for irradiation). Each harvested replicate was split between the control and irradiated sets so that there were paired unirradiated and to-be-irradiated replicates originating from the same initial recovered culture inoculated from frozen stock. The bottles (unirradiated controls) were put on ice and kept in darkness. The culture tubes were put on ice in the custom γ-irradiator culture tube rack for transport and subsequent irradiation. Ice was packed tightly into the custom rack to capacity after inserting culture tubes, adding ∼250 mL tap water, shaking, and tightly packing with ice again. A crevice was poked into the center of the packed ice using a 15 mL conical tube and filled with 4 dry ice pellets. These conditions cool internal liquid culture temperature to < 4°C in about 10 min. The cap was placed on the rack, and the rack was set inside a Styrofoam box filled with ice for transport to the irradiator. Time from placement on ice until start of irradiation was about 30 min, including transport time.

#### Gamma-irradiation experiment

Samples in the γ-irradiator culture tube rack were placed in the sample chamber of a GC-220E ^60^Co γ-irradiator (dose rate = 55.18 Gy/min) and irradiated for 2 hours, receiving a total dose of 6.7 kGy. After irradiation, the γ-irradiator culture tube rack was refilled with fresh ice and transported (30 min) before protein extraction. Maintaining cultures near 0°C for 30 min before, throughout 2-h irradiation, and 30 min after ensured that proteomic changes were primarily due to oxidation-induced damage and not regulation of *de novo* gene expression. This is because of the deleterious effects of irradiation and cold on transcription and translation combined with the short timescale of our experiment.

Protein carbonylation leads to loss of protein function and lack of detection by shotgun proteomics due to aggregation, self-cleavage, and proteolysis (*12*). Aggregation accounts for ∼95% of carbonylated proteins in *E. coli* (*50*), affecting solubility and detection by proteomics (*51*). Proline carbonylation can lead to non-enzymatic self-cleavage at the peptide backbone via the proline oxidation pathway (*13*). Low temperature slows all enzymatic activity, but proteases targeting carbonylated proteins retain at least some rate at low temperature, such as the *E. coli* Lon protease functioning at 16°C (*52*).

Regulation of transcription and translation in *D. radiodurans* in response to irradiation does not substantially occur until return to permissive growth conditions. Previous irradiation experiments without low temperature show that radiation-responsive promoters in *D. radiodurans* do not activate transcription until after irradiation stops, and peak transcriptional rates occur 1 to 2 h into recovery in fresh medium at optimal temperature (*53*); similar trends hold transcriptome-wide (*40*). RNA samples taken mid-irradiation, without post-irradiation recovery, result in nearly 50% fewer reads originating from mRNA relative to pre-irradiation or during recovery, and nearly all individual transcripts decrease (*54*). Similar to transcription, *D. radiodurans* proteins expressed in response to irradiation without cold temperature reach their peak translation rate 0.5 to 1 h after transfer to fresh media under optimal temperature (*49*).

Transcription and translation both substantially slow or halt under cold temperatures in *E.coli* and *D. radiodurans*. *D. radiodurans* and *E. coli* RNA polymerases slow 30-fold transitioning from 37°C to 0°C (*55*). *E. coli* translational elongation is slowed 3-fold transitioning from 37°C to 25°C (*56*) and >10-fold from 37°C to 10°C (*57*). *E. coli* protein synthesis slows to a halt after 30 min at 0°C because translation cannot initiate (*58*). In *D. radiodurans*, cold shock without irradiation only influences ∼5% of expressed proteins after 3 h at 20°C (*59*), and translation occurs only very little at 0°C (*60*). Taken together, proteomic changes observed in our experiments must be due to oxidation-induced damage leading to a combination of aggregation and degradation as opposed to *de novo* protein synthesis.

Small volumes (110 μL) of each irradiated and unirradiated replicate were reserved in 1.5 mL microfuge tubes and held on ice in dark while protein extraction was initiated (see below).

Serial dilutions of each irradiated and unirradiated replicate were made: undiluted, 1:1,000, and 1:10,000 dilutions in LB for *E. coli*; and undiluted, 1:10,000, and 1:100,000 dilutions in TGY for *D. radiodurans*. We chose different dilution factors for each species to facilitate colony counting in anticipation of drastically different irradiation survival rates. 100 μL of each dilution was spread on pre-warmed solid LB medium or TGY medium plates for *E. coli* and *D. radiodurans*, respectively. *E. coli* plates were grown overnight at 37°C in dark, and *D. radiodurans* plates were grown for 3 days at 30°C in dark. Colony forming units (CFUs) were counted at the end of each growth period for each plate, and the irradiation survival rate (Fig. S2A) was computed for each replicate as the ratio of CFUs on plates with irradiated samples to CFUs on plates with unirradiated samples. Due to our irradiation treatment, no *E. coli* colonies formed even from undiluted irradiated cultures, and 55-70% survival of CFUs was observed for *D. radiodurans* (Fig. S2A).

#### Protein extraction and derivatization protocol

The protein extraction protocol was performed as soon as possible after irradiation and after reserving culture volumes to make serial dilutions for survival assay; this was in an effort to retain as much oxidized protein as possible.

1. Lysis buffer was warmed to room temperature on a nutator to dissolve solutes with occasional vortexing.
2. Remaining volume of each sample was poured evenly into three 50 mL conical tubes and centrifuged at 4,000 g for 8 min at 4°C.
3. Pellets were combined for each sample into one tube and washed using 25 mL harvest buffer. Samples were spun down once more and supernatant removed. Samples were kept in the same 50 mL conical tubes for lysis.
4. Cells were resuspended in 276 μL lysis buffer.
5. To each sample, 46 μL of 7X protease inhibitor cocktail was added.
6. Cells were broken with 3 freeze/thaw cycles, each cycle consisting of 15 min on dry ice and thawing at room temperature for 15 min, vortexing between thawing and re-freezing.
7. Cells were sonicated in 4°C cold room using a digital Fisher Scientific 550 Sonic Dismembrator (amplitude 20%, 30 cycles of 8 s pulse, 10 s off). The base of each tube was held on ice before, during, and after sonication to cool the sample.
8. The samples were transferred to microfuge tubes and subsequently centrifuged at 8,000 g at 4°C for 10 min.
9. The supernatant for each sample containing soluble proteins was transferred to a 2 mL microfuge tube.
10. Proteins were derivatized with 300 μL derivitization buffer for 1 h in the dark on a rotating mixer at 4°C. The amine of DNPH (-NH-NH2) reacts specifically with the carbonyl functional groups in aldehydes and ketones to form corresponding 2,4-DNP-hydrazones.
11. To stabilize hydrazones, 300 μL of hydrazone stabilization buffer was added for 1 h in the dark on a rotating mixer at 4°C.
12. Protein samples were precipitated with 225 μL protein precipitation buffer for 3 h in the dark on a rotating mixer at 4°C and then centrifuged for 15 min at 10,000 g at 4°C.
13. Protein resolubilization buffer was warmed in a 55°C water bath briefly with occasional vortexing until solutes dissolved.
14. Protein pellets were washed twice by resuspending in 1 mL wash buffer, centrifuging for 15 min at 10,000 g at 4°C, and removing supernatant in order to remove excess DNPH.
15. Samples were resuspended in 110 μL protein resolubilization buffer at 37°C for 40 min with shaking in an Eppendorf Thermomixer with occasional manual mixing with clean pipette tips to help break up pellets.
16. To estimate protein concentrations and determine the amount of trypsin needed for protein digestion, 10 μL of each sample was set aside in cryotubes and stored at -80°C for later quantification by Bradford assay and visual confirmation by SDS-PAGE.
17. Remaining samples were also stored at -80°C.

#### Bradford assay protocol

Measurement of protein concentration was performed as follows:

1. Reserved 10 μL protein samples were thawed on ice while Bradford reagent (Coomassie from Pierce™ Coomassie (Bradford) Protein Assay Kit, ThermoFisher 23200) was warmed to room temperature.
2. Thawed protein samples were diluted 1:50 (1 μL reserve, 49 μL distilled water) and 1:100 (25 μL of 1:50 dilution, 25 μL distilled water) in fresh microfuge tubes.
3. 5 μL of each dilution was loaded into a separate well of clear-bottom 96-well plate.
4. 250 μL Bradford reagent was loaded into in each well.
5. Samples stained at room temperature for 10 min.
6. Bovine Serium Albumin (BSA) standards from Bradford assay kit were loaded into empty wells in the 96-well plate.
7. Samples on plate were loaded into a plate reader and their absorbance at 600 nm measured in 3 technical replicates.
8. Second-order regression function was fitted to BSA standard concentration data and used to estimate protein sample concentrations, correcting for dilution factors. Final concentrations for protein samples were also projected based on volume changes due to reducing, alkylation, and pH adjustment steps as part of the trypsinization protocol (see below).

#### SDS-PAGE protocol

Protein sample quality was confirmed visually as follows:

1. A volume corresponding to 40 μg of each protein sample reserve thawed for the Bradford assay, or as close as possible not exceeding 10 μL, was transfered to a separate fresh microfuge tube.
2. Total volume in these tubes was adjusted to 10 μL with distilled water.
3. 5 μL 3X SDS-PAGE loading buffer was added to each sample, briefly vortex, and boiled at 100°C for 15 min.
4. While boiling protein samples, Novex WedgeWell 14% Tris-Glycine Gel cartridges were prepared by rinsing with tap water, pulling out comb, rinsing again with tap water, removing strip on cartridge to expose bottom of gel, and inserting cartridge into gel dock. Inner reservoir of gel dock was filled completely and outer reservoir filled to about 2 cm with Tris-Glycine buffer, to just above the exposed gel where the strip was.
5. Protein samples were removed from 100°C, briefly vortexed, and centrifuged at full speed for 10 min.
6. Supernatants were loaded from sample tubes into gel wells, and 6 μL of Color Prestained Protein Standard, Broad Range (11-245 kDa) (NEB P7712S) was loaded into an empty well.
7. Gel was run at 100 V for 1.5 h and stopped when dye front neared the outer reservoir buffer.
8. Gel was carefully transferred from cartridge into an empty staining dish.
9. 125 mL stain solution was heated in the microwave for 20 seconds and then poured into the staining dish away from the gel until the gel was just completely covered.
10. Staining dish was covered with plastic wrap, and gel was stained on nutator with gentle agitation for 5 min. After staining, excess stain was poured back into stock bottle for reuse.
11. Gel was destained by pouring about 300 mL of destain solution into dish away from gel and gently agitating on nutator for 15 min. Destain solution was replaced with about 400 mL of fresh destain solution and gently agitated on nutator overnight (18 to 20 h).
12. Gel was imaged in Bio-Rad ChemiDoc MP using white light.

#### Trypsinization and speed-drying protocol

1. Derivatized protein samples stored at -80°C were thawed on ice. For all subsequent steps until overnight digestion, sample and trypsin tubes were kept on ice.
2. Protein samples were reduced with 12.2 μL freshly prepared reducing buffer for 30 min at 60°C in Eppendorf Thermomixer.
3. Protein samples were alkylated with 40.7 μL freshly prepared alkylation buffer for 30 min at 25°C in Eppendorf Thermomixer.
4. Sample pH was checked using 2 μL sample on pH paper. *D. radiodurans* samples were near pH 7.0 and *E. coli* near pH 6.0. All samples were adjusted to pH 8.0 by adding 1 μL and 2 μL of 10 M NaOH to *D. radiodurans* and *E. coli* samples, respectively. This pH adjustment step is critical for optimal trypsin activity.
5. For each sample, a volume corresponding to a protein mass of 300 μg, or as close as possible without going over 300 μg or 150 μL, was transferred to a fresh microfuge tube.
6. Sample tubes containing less than 150 μL of sample were raised to a total volume to 150 μL by adding an appropriate volume of BioRad ReadyPrep™ Proteomics Grade Water. For the remaining trypsinization steps, a protein mass of 300 μg was assumed for all samples and a trypsin-to-protein ratio of 1:50 (w/w) was used.
7. The necessary number of lyophilized trypsin vials (Trypsin from porcine pancreas Proteomics Grade, BioReagent, Dimethylated, Sigma T6567-5X20UG) were used for all samples being digested; each vial contains 20 μg trypsin. Caps were removed from the vials.
8. 20 μL trypsin solubilization reagent (1 mM HCl) was added to each trypsin vial to give 1 μg/μl reconstituted trypsin.
9. To each protein sample, 144 μL of trypsin reaction buffer was added.
10. To each protein sample, 6 μl of the reconstituted trypsin was added for a total reaction volume of 300 μL. Sample was mixed well by vortexing.
11. Samples were digested at 37°C overnight (16 h).
12. Digested samples were briefly spun down for condensation that formed in tube caps and transferred to fresh 1.5 mL conical polypropylene tubes with screw caps (without skirts) to be dried immediately. Screw caps were left off for drying.
13. Samples were dried using a GeneVac EZ-2 Plus HCl-compatible (model EZ-2.3). The temperature was set as low as possible with the lamp off. The Low BP Mixture setting was used, and the run time was set to 2 h and 45 min to final stage and a final stage time of 15 min.
14. Dried samples were unloaded, closed with screw caps, and put on ice.
15. Dried samples were transported to mass spectrometer on dry ice.

#### Liquid chromatography tandem mass spectrometry (LC-MS/MS)

Protein identification and quantification was performed using a label-free strategy on a UHPLC-HRMS platform composed of an eksigent 2D liquid chromatograph and an AB SCIEX TripleTOF™ 5600. Peptides were separated on a 25 cm C18 column (Acclaim pepmap100, 3 µm, Dionex) by a linear acetonitrile (ACN) gradient (5–35% (v/v), in 15 or 120 min for short and long runs, respectively) in water containing 0.1% (v/v) formic acid at a flow rate of 300 nl/min. In order to reach high retention stability, which is a requirement for label-free quantification, the column was equilibrated with a 10x volume of 5% ACN before each injection. Eluant was sprayed using the Nanospray Source into the TripleTOF™ 5600. Mass spectra (MS) were acquired across 400–1500 m/z in high resolution mode (resolution >35000) with 500 ms accumulation time. The instrument was operated in DDA (data dependent acquisition) mode and MS/MS were acquired across 100–1800 m/z. For short runs, precursor selection parameters were: intensity threshold 400 cps, 20 precursors maximum per cycle, 100 ms accumulation time, 10 s exclusion after one spectrum. For long runs, precursor selection parameters were: intensity threshold 200 cps, 50 precursors maximum per cycle, 50 ms accumulation time, 30 s exclusion after one spectrum. A long run procedure was used to acquire quantitative data, and a duty cycle of 3 sec per cycle was used to ensure that high quality extracted ion chromatograms (XIC) could be obtained.

#### Modified peptide identification

Protein searches for mass spectra obtained on the Triple-TOF 5600 LC-MS/MS were performed against a local copy of the *D. radiodurans* R1 (UP000002524) and *E. coli* K-12 MG1655 (UP000000625) database (retrieved from UniProt on June 26, 2017; 3085 and 4307 proteins respectively) using ProteinPilot^TM^ 5.0.1 Revision 4895 (Paragon^TM^ Algorithm 5.0.1.0, 4874). One missed internal tryptic cleavage site per peptide was accounted for in the search parameters. Mass tolerance was set to 15 ppm in MS and 0.05 Da in MS/MS. In addition to the standard biological modifications set including a differential amino acid mass shift for oxidized methionine (+15.99 Da), custom modifications accounting for DNPH derivatization adducts (Fig. S1), both with and without a typical neutral loss (noNL), were added to the data dictionary and parameter translation files (File S3 and File S4). These custom modifications included:

- Arginine → DNP-glutamic-5-semialdehyde (+136.97 Da)
- Lysine → DNP-allysine (+179.00 Da)
- Proline → DNP-glutamic-5-semialdehyde (DNP-Glu5A) (+196.02 Da)
- Proline → DNP-pyroglutamic acid (DNP-PCA) (+194.01 Da)
- Threonine → DNP-2-amino-3-oxobutanoic acid (DNP-AKB) (+178.01 Da)

The Paragon^TM^ Algorithm was run to identify peptides from each replicate sample with the following parameter settings:

- Sample Type: Identification
- Cys Alkylation: Iodoacetamide
- Digestion: Trypsin
- Instrument: TripleTOF 5600
- Special Factors: Urea denaturation; DNPH derivatization; DNPH derivatization noNL
- Species: *Deinococcus radiodurans* (OR *Escherichia coli*)
- ID Focus: Biological modifications
- Database: Deinococcus_radiodurans.fasta (OR Escherichia_coli.fasta)
- Search Effort: Thorough ID
- Detected Protein Threshold (Unused ProtScore (Conf)) >: 0.05 (10.0%)
- Run False Discovery Rate Analysis: (yes)

High confidence peptides were selected using a 99% confidence threshold for proteins identified by at least 2 peptides (Table S1). After identifying peptides from all sample replicates, carbonyl modifications were taken as those residues exhibiting one of the DNPH modifications described above or any of the following default modifications included in the data dictionary file that represent the direct products of RKPT side chain oxidation by ROS:

- Arginine → Glutamic 5-semialdehyde (-43.05 Da)
- Lysine → Allysine (-1.03 Da)
- Lysine → 2-aminoadipic acid (double oxidation of lysine) (+14.96 Da)
- Proline → Glutamic 5-semialdehyde (+15.99 Da)
- Proline → Pyroglutamic acid (+13.98 Da)
- Threonine → 2-amino-3-oxobutanoic acid (-2.02 Da)

The false discovery rate (FDR) was calculated at the peptide level for all experimental runs using ProteinPilot; this rate was estimated to be lower than 1% for both *D. radiodurans* and *coli*.

#### Estimation of total CS in samples by saturation curve fitting

Saturation curve fitting was used to estimate the total number of CS in shotgun redox proteomic samples, a technique that has been demonstrated for phospho-proteomics when pooling data across many studies (*61*). To this end, we identified the number of redundant and non-redundant CS within each replicate experiment for each organism and computed the cumulative totals with the addition of each successive replicate (Fig. S2B). Exponential saturation functions were fit by minimization of the sum of squared errors with experimentally-determined data points. The fit exponential functions asymptotically approach theoretical maxima, which represent the estimated total CS in our samples. The estimated percentage of sampling coverage is then taken as the ratio of total measured non-redundant CS to total estimated non-redundant CS for each species. Fit curves that remain very linear suggest very low sampling coverage, and curves that saturate suggest near-complete sampling coverage of unique sites. The more replicate experiments available, the higher the confidence in such estimates.

#### Protein quantification

For quantification, the quant application of PeakView was used to calculate XIC for all peptides identified with a confidence > 0.99 using ProteinPilot™. A retention time window of 2 min and a mass tolerance of 0.015 m/z were used. The area under the curve was exported in MarkerView™, in which they were normalized based on the summed area of the entire run.

MarkerView™ enabled an average intensity for irradiated and unirradiated conditions to be calculated, as well as the significance of the difference between conditions based on a paired t-test. Quantified proteins were kept with a p-value < 0.1 and with at least one peptide quantified with a p-value < 0.1. In order to be considered as significantly changed in relative abundance, proteins had to meet two criteria: 1) mean relative abundance fold change within a cut-off of > 2-fold (for increased) or < 0.5-fold (for decreased) in the irradiated samples relative to the controls, and 2) the paired t-test p-value had to be < 0.05.

#### Amino acid prevalence analysis

For RKPT carbonylation prevalence, only RKPT on peptides bearing at ≥ 1 CS were included. For pre-irradiation prevalence and change in prevalence after irradiation, all amino acid residues in peptides identified as described above were included. Occurrence of each amino acid was counted within the respective set of included peptides, and relative proportion was computed as the percentage of the total number of residues. Statistical significance of differences in relative proportions was determined using the z-test for independent proportions. The significance threshold was taken as results with p-value < 0.01.

##### Functional classification analysis of proteomics data

The aGOtool web server (*25*) was used to perform Gene Ontology (GO) (*62*) biological process term enrichment analysis among groups of proteins of interest. These groups included proteins for which carbonyl sites were identified or for which relative abundance changes after irradiation were > 2-fold or < 0.5-fold in *E. coli* or *D. radiodurans*. To correct for protein abundances, the background mean normalized abundance pre-irradiation was used for all detected proteins. A p-value cutoff of < 0.05 was used for reporting over- and underrepresented GO terms.

##### Selection of positive and negative experimentally-identified CS for computational analysis

From all high-confidence modified peptide calls, taken as described above, all residues bearing RKPT side chain carbonyls or derivatized labels for carbonyls were assigned as positive CS. Unmodified RKPT on this same set of peptides were counted as negative CS. All positive and negative CS and the peptide sequences with modifications used to assign them are listed in Table S7.

##### Protein structure derivation

###### Target proteins

The target proteins consisted of all genomically-encoded *D. radiodurans* R1 proteins.

Amino acid sequences for target proteins (3184 proteins) were taken from the translated *D. radiodurans* R1 genome (*63*) as annotated by the RAST server (*64*) as of May 4, 2016 and reconciled against their respective UniProt entries when available (*65*). An index of all target proteins and statistics for their structures is listed in Table S3.

###### Curation of experimentally solved structures

Experimentally solved structures were curated from the PDB (*66*), when available, for all D *radiodurans* target proteins. The best PDB structure from alternative available structures for each protein was chosen based on the best resolution structure with highest coverage of the full-length protein sequence. The chosen structures were edited to remove all ligands and all but the single best-representative chain for each protein. This resulted in 71 *D. radiodurans* proteins covered by crystal structures.

###### Structure modeling

Structures were modeled for all *D. radiodurans* target proteins using 5 methods, as permitted by the technical constraints of each method. These methods included I-TASSER (*23*), QUARK (*34*), modeler and scwrl algorithms as implemented on the ProtMod server (http://protmod.godziklab.org/protmod-cgi/protModHome.pl) (*67, 68*), and EVfold (*69*).

Developer-recommended protein length limits and high quality homologous template availability, as summarized in Table S4, determined which methods could be applied for each target protein. For targets modeled by I-TASSER, the LOMETS (*70*) software was used to determine the number of high quality homologous templates available and categorize each target as easy (> 10 templates), medium (1 to 10 templates), or hard (0 templates) to model. For targets modeled using the ProtMod server, the FFAS-3D (*71*) software was used to select the single most appropriate homologous template. The top ranking of alternative models from each method was selected for further analysis as determined by scoring metrics particular to each method.

Each method was run using default parameters as noted in their respective publications or as included as default settings for the web server implementations of each method as available.

Other than for training set proteins, which were curated from the PDB and modeled comprehensively as described above, *E. coli* target protein structures were taken as modeled previously (*72*) using I-TASSER and QUARK across the whole proteome. Given that either I-

TASSER or QUARK was the top performing modeling method for the majority of proteins in the *D. radiodurans* proteome and given the high coverage of the *E. coli* proteome by experimentally solved structures, comprehensive modeling of *E. coli* proteins using modeler, scwrl, and EVfold was seen as unlikely to yield higher quality structures for *E. coli* proteins.

#### Quality evaluations and selection of best-representative structures

The best PDB structure and model from each method were all evaluated for quality with respect to 8 metrics (Table S5): target sequence coverage, computable surface using UCSF Chimera (*73*), 3 separate energy scores (*74–76*), estimated TM-score (*77*), percentage of residues in favored positions with respect to a Ramachandran plot (*78*), and an overall conformation score (*78*). Two criteria were absolutely required to be included in downstream analyses: >90% target sequence coverage and computable surface. This is because high structural coverage for each protein was seen as critical for comprehensive evaluation of possible CS and because several molecular properties relied on protein surface computation. The criteria used for favorable quality evaluation with respect to the other metrics were derived from publications of respective methods as noted in Table S5. The best-representative for each protein was chosen based on which structure satisfied not only the two absolutely required criteria but also the highest total number of satisfied criteria. Ties for total highest number of satisfied criteria were broken by comparing the means of maximum-normalized metrics for the tying structures. Only the best-representative structure for each protein was used in subsequent structural analysis and feature computation for machine learning.

##### Feature engineering

###### Selection of molecular properties

The published literature on protein oxidation and modification was reviewed in search of protein molecular properties previously hypothesized or tested for contribution to susceptibility or robustness to damage by ROS. Rate constants for reaction with hydroxyl radical were included with respect to each type of amino acid (*79*). As a proxy for reaction rate constants with all ROS that might lead to carbonyl formation on RKPT, the relative reactivity of each of these four amino acids was derived based on the relative proportion of experimentally-measured CS distributed across the four amino acids and relative proportion of non-oxidized occurrence distributed across the four amino acids, an approached described previously (*26*).

The more exposed a residue is, the higher probability of contact with any molecules in the solvent (*80–82*). Therefore solvent accessible surface area and depth (i.e. minimum distance from the protein surface) were computed using the surface computation and distance measurement tools in UCSF Chimera (*73*). Similarly, residues that form secondary structure often have less exposed side chains; the UCSF Chimera implementation of the DSSP algorithm (*83*) was used to assign secondary structure. Molecular density can also be expected to sterically interfere with small molecule interaction; circular variance (*84*) was computed as a proxy for molecular density. Number of contacts, another measure of density, was computed based on interatomic distances using UCSF Chimera. Protein-protein interactions can also be expected to interfere with small molecule interaction, and DISOPRED (*85*) and SPPIDER (*86*) were used to assess residues likely to take part in protein binding sites.

Correlation of dehydration tolerance with both disorder and hydrophobicity has been demonstrated previously (*87*); disorder was computed using DISOPRED (*85*) and hydrophobicity using the Kyte-Doolittle scale (*88*).

Electrostatic charge is expected to impact protein interaction with charged molecules (*89*), such as hydroxyl and superoxide radicals; therefore, canonical formal charges for amino acids were included.

Metal binding sites for Fe and Cu cations can cause residues nearby to undergo metal-catalyzed oxidation (*29*), but metalloproteins coordinating Mn cations are involved in sequestering ROS (*90*). Experimentally-characterized metal binding sites were curated from UniProt (*65*) and mapped to protein structures included in this study, and to expand the very sparse available experimental data, FINDSITE-metal (*91*) was used to predict metal binding sites from structural homology with homologous templates found using LOMETS (*70*).

Surface methionines and cysteines can serve to protect other nearby residues from oxidative damage through their own reversible oxidation (*29*); these amino acids were included as found on protein surfaces defined using UCSF Chimera.

To further broaden the search space of potentially useful molecular features a database of hundreds of amino acid properties (*92*), as summarized by 5 factors called AAMetrics (*93*), was included in this study. These factors can be conceptually summarized as corresponding to electrostatic charge, propensity to form secondary structure, molecular volume, codon diversity, and polarity.

#### Features from parameterization of molecular properties

In the conversion of molecular properties to features, several parameters for computation were varied:

Scale: In addition to computing each molecular property with respect to the atomic location of possible carbonyl formation on each RKPT (i.e. residue scale), each property was also computed with respect to all residues within a defined radius (5Å, 8Å, 10Å, and 12Å) and with respect to the entire protein to account for the global scale. The radii used for local-scale feature computation were selected based on effective protein contact distances previously determined in published studies (*84, 89, 94, 95*). The residue scale is not applicable to surface cysteine and methionines, contacts, RKPT contacts, or negative formal charge because of the type of amino acids under consideration or due to logical exclusions.

Summary statistic: As a summary statistic of each molecular property, the sum and mean of component values were computed. At the residue scale, the sum and mean are equivalent because there is only one component value. The mean summary statistic is not applicable to metal binding site features, contacts, or circular variance.

Grouping: When applicable, molecular properties were converted to features as a composite and as separate features. Applicable properties included surface cysteines and methionines, canonical charge, RKPT contacts, secondary structure, and metal binding sites. For example, separate features were computed with respect to binding sites for each elemental type of metal cation, and also, features were computed with respect to all metal cation binding sites irrespective of elemental type.

Data type: Some molecular properties may be treated as either continuous or binary variables; each option was used to yield separate features with respect to such properties.

Applicable properties included protein disorder and protein binding sites as output by DISOPRED, which outputs these properties both as a continuous probability score for each residue and as a binary classification of disordered or not.

#### Features from sequence alignment

In addition to structure-based features, local amino acid sequence homology was used to generate features to represent positive and negative CS. We defined a local neighborhood of 21 residues centered on each RKPT covered by carbonyl-bearing peptides in our proteomic data.

All-by-all semi-global alignments of the non-redundant subset of these sequences was performed, setting the central RKPT of each 21-mer as anchor points to seed each alignment. The resultant alignment score matrix had the same dimensionality as the number of non-redundant RKPT sites in our CS positive and negative data (979); therefore, feature reduction was performed during the machine learning phase (see below) by principal component analysis (PCA).

##### Machine learning

###### Training data

The training set consisted of 397 molecular-property-derived features computed using best-representative protein structures and 979 sequence-alignment-derived features corresponding to experimentally-measured positive and negative CS from *E. coli* and *D. radiodurans* proteomes. This dataset includes 869 non-unique positives and 2777 non-unique negative CS in 153 unique proteins. The full training data set is included in Table S7. All features in the training set were standardized by mean centering at 0 and scaling to unit variance using the scikit-learn StandardScaler function. The same scaling factors from the training data were used for standardizing the test data.

###### Logistic regression algorithm

We used 64-bit Python version 3.6.2 with Scikit-learn version 0.19.1 for all machine learning in this study. To model the probability of RKPT carbonylation, we used a logistic regression estimator with stochastic gradient descent training, a ‘log’ loss function, and elastic net regularization that linearly combines L1 and L2 penalties. The elastic net mixing parameter for L1 and L2 penalties was set to 0.5 and the learning rate to 0.1 to balance utilization of informative features with elimination of extraneous features. Fitting the logistic regression model with these parameters will drop many coefficients of uninformative features to zero. Balanced class weighting was used to account for the class imbalance in training data such that weights for each class *i* (positive or negative) are inversely proportional to the class size *n* for data sample 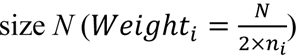

Logistic regression models were fit separately to structure-based and sequence-based features. While the regularization strategy described above was sufficient to reduce the number of features in the structure-based model, the great number of sequence-based features and their relatively low individual predictive power required an additional feature reduction step. To this end, we performed PCA on the sequence alignment score matrix, using just the top principal components collectively accounting for 95% of data variability as features for the sequence-based model.

We combined the sequence-based and structure-based logistic regression models into a final stacked model. Input to the stacked model consists of features with non-zero coefficients from the structure-based model and the PCA-derived features from the sequence-based model along with two meta features representing the individual predicted probabilities from each model. The stacked model was fitted with L2 regularization (but not L1) to attain a stable solution that does not exclude any of the informative features from the initial two models. The magnitude and sign of the fit model coefficients quantify the relative contribution of each feature to overall probability estimation. This implies that molecular properties underlying contributing features are predictive of oxidation probability but does not necessarily imply that molecular properties represented by features with zero coefficients cannot influence oxidation by ROS *in vivo*, simply that our data do not support the predictive power of those features.

###### Leave-1-out validation strategy

To assess generalization of predictive performance of our predictive framework, comprehensive leave-1-out cross validation was performed. Because the training data contains duplicate data points due to coverage of particular RKPT by multiple measured peptides, we implemented a variant of standard leave-1-out validation in which all duplicates of a data point (positive or negative) are treated as one data unit, i.e. validation is performed on them together in one leave-1-out iteration. Also, because our full set of sequence-based features contains information derived from local sequence surrounding every RKPT in the training data, we excluded sequence-features corresponding to alignment scores against the held-out data for each leave-1-out iteration before performing PCA as described above.

##### Multi-scale model validation

###### Residue-scale validation

We used Receiver Operating Characteristic (ROC) curve analysis to validate the overall performance of our machine learning framework for protein carbonylation site probability estimation. Area under the ROC curve (AUC) is a common, robust performance metric for machine learning predictors and is not affected by data imbalance, which is especially important for predicting protein CS because redox proteomic techniques yield a relatively much smaller number of positives than negatives in peptide calls. The calculation of the area under the ROC curve was weighted by class sizes using the same weighting factor described above.

Furthermore, because of the stochastic oxidation of RKPT observed in the training data (i.e. the same site can appear oxidized on one peptide but not oxidized on another), an AUC of 1.0 (a perfect predictor score) is not guaranteed to be achievable. Therefore, we computed the theoretical maximum ROC curve and corresponding AUC for our data (Fig. 5A lower left) using empirically-derived carbonylation probabilities. All reported AUCs were normalized to this theoretical maximum AUC.

We also performed a randomization test in which the initial structure-based and sequence-based features were first shuffled before running the entire predictive framework. This randomization test gives us a sense of the non-random significance of predictive features in our structure-based, sequence-based, and stacked models. We also compared our model performance to that of the Carbonylated Site and Protein Detection (CSPD) model (*17*). To test CSPD performance on our data, we used the CSPD web server and input all full-length protein sequences from our experimental dataset. CSPD output is a binary classification of carbonylation sites, rather than a probability estimator. ROC analysis was performed on these predictions as well.

###### Protein-scale validation by carbonyl site enrichment correlation

To validate our protein carbonyl prediction framework at the whole-protein scale, we computed a predicted enrichment score for each protein as the sum of the predicted probabilities for sites included in the carbonylated peptide data normalized by the length of the protein regions covered by those peptides. Similarly, an enrichment score was computed based solely on the carbonylation state of RKPT residues in the experimentally measured carbonylated peptides.

These scores appear in the x- and y-axes of Fig. 5B. Predicted oxidation enrichment was validated against all 90 *E. coli* and 63 *D. radiodurans* proteins detected with at least one CS by redox proteomics and coverage by our 3D protein structures. Spearman rank correlation between predicted and experimental enrichment scores was computed for proteins from each species, and 95% confidence intervals were determined for a fitted linear regression (Fig. 5B).

##### Interspecies proteome-wide prediction

###### Data sets for prediction

After using our framework to train a model on the entire training set, we applied our predictor prospectively to compare protein carbonylation propensity across the full proteomes of *coli* (4,057 proteins) and *D. radiodurans* (3,031 proteins), containing 227,326 and 184,149 total RKPT, respectively. The full proteome data sets for *E. coli* and *D. radiodurans* are included in Table S8 and Table S9.

###### Ortholog and isozyme mapping

In order to directly compare individual proteins between *E. coli* and *D. radiodurans*, pairs of proteins were mapped by shared function between these species. This was done in three steps. First, functional annotation mapping between species was performed using the RAST server (*96*) for annotation and the ModelSEED server (*97*) for mapping. This approach has the ability to map orthologs between species as well as non-homologous isozymes that only share function.

Second, likely orthologous pairs were mapped between species by bi-directional BLAST (*98*) in which mutual top hits between species are identified, and for cases where proteins do not have mutual top hits by bi-directional BLAST, simply the top hit in the other species is identified.

Tying top hits with equal E-values and alignment scores were retained at this step. Finally, manual curation of the mapped interspecies pairs was performed, reconciling functional annotation of *E. coli* proteins in EcoCyc (*99*) and *D. radiodurans* protein annotations in BioCyc (*100*). During the curation process, proteins from one species mapping to multiple proteins in the other species were reduced to mapping to as few proteins from the other species as possible based on a combination of functional annotation specificity and any fine differences in sequence alignments. As a result, 1170 *D. radiodurans* proteins were mapped to 1110 *E. coli* proteins for a total of 1300 interspecies pairs that also have structural coverage in our data (Fig. S4).

###### Pairwise comparison of predicted vulnerability to oxidation

To compare protein-scale predicted oxidation susceptibility between proteins from each species, we computed an enrichment score for each protein equal to the number of RKPT with probability > 0.5 for carbonylation normalized by the protein length. These scores appear in the x- and y-axes of Fig. 7.

To quantify the relative protein-intrinsic vulnerability to oxidation between interspecies pairs, we take the perpendicular distance of the point representing the pair to the y=x diagonal reference line. Proteins less intrinsically vulnerable to oxidation in one organism lie a greater distance to one side of y=x. To quantify the combined protein-intrinsic and extrinsic differences between interspecies pairs, we take the perpendicular distance of the point representing the pair to the y=x/3.78 reference line. The 3.78 factor represents the ratio of protein carbonyls generated *in vivo* in *E. coli* compared to *D. radiodurans* after exposure to 7kGy γ-radiation (ratio = 3.86), minus the contribution from protein-intrinsic factors alone. This latter contribution from protein-intrinsic factors comes from the ratio of protein carbonyls measured in *E. coli* lysate compared to *D. radiodurans* lysate after dialysis to remove ions (ratio = 1.08). Proteins less susceptible in *D. radiodurans* than in *E. coli* when accounting for intrinsic and extrinsic factors lie a greater distance above y=x/3.78.

Extreme outliers of the interspecies comparison are expected to be involved in cellular resistance to oxidative stress, if indeed *D. radiodurans* proteins have evolved intrinsic properties to protect them from oxidation. We define a boundary to highlight extreme outliers of the all-pair distribution based on a minimum-fitted ellipse that encompasses all points within 3 standard deviations of the mean coordinates and within 3 standard deviation of the mean distance from y=x. Outliers to this boundary account for 8% of all interspecies mapped protein pairs.

---

**Fig. S1.**
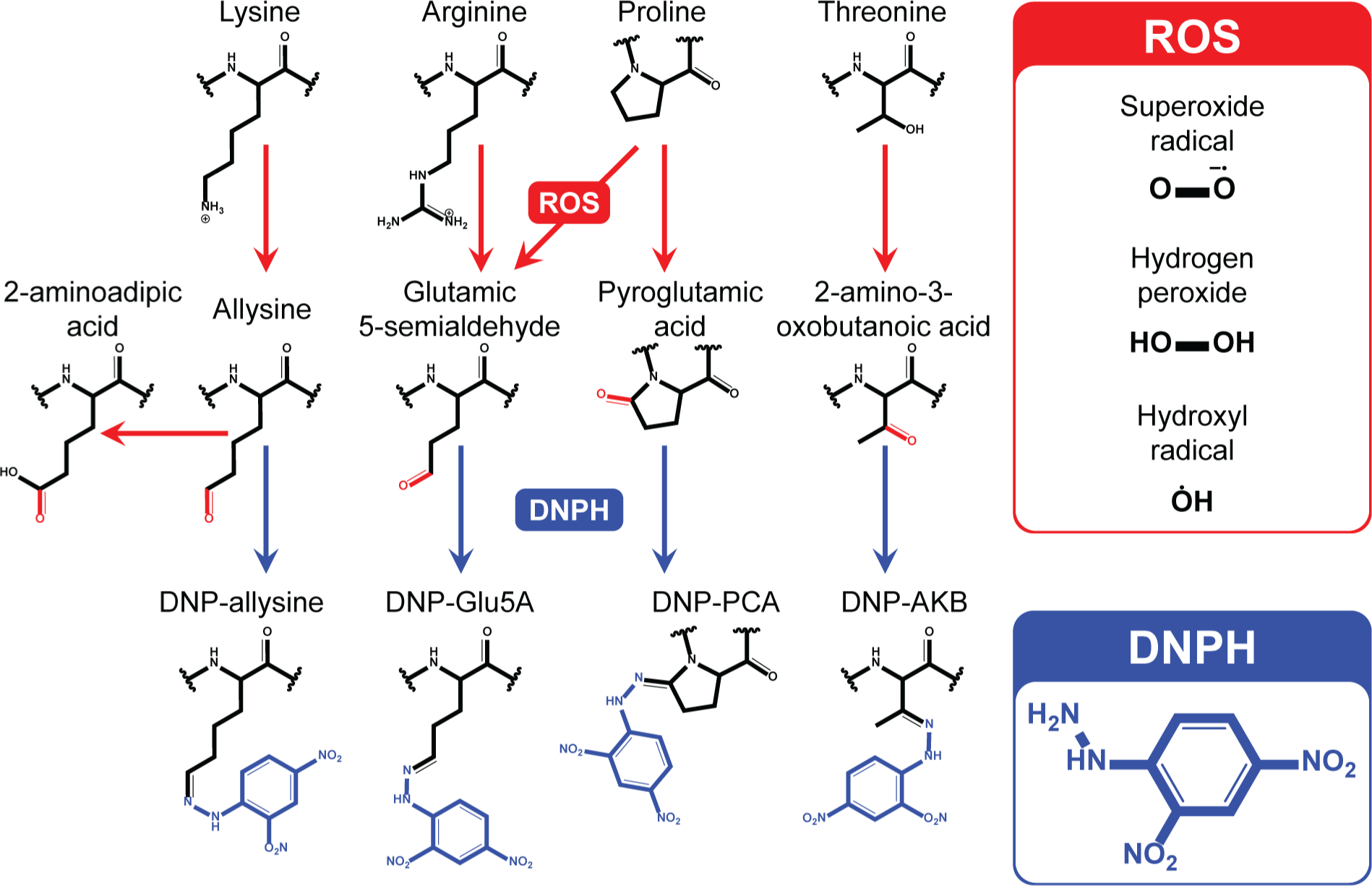
Carbonyl formation pathways for RKPT reacting with ROS and adducts derivatized by DNPH, related to Fig. 2A and 3A.

**Fig. S2.**
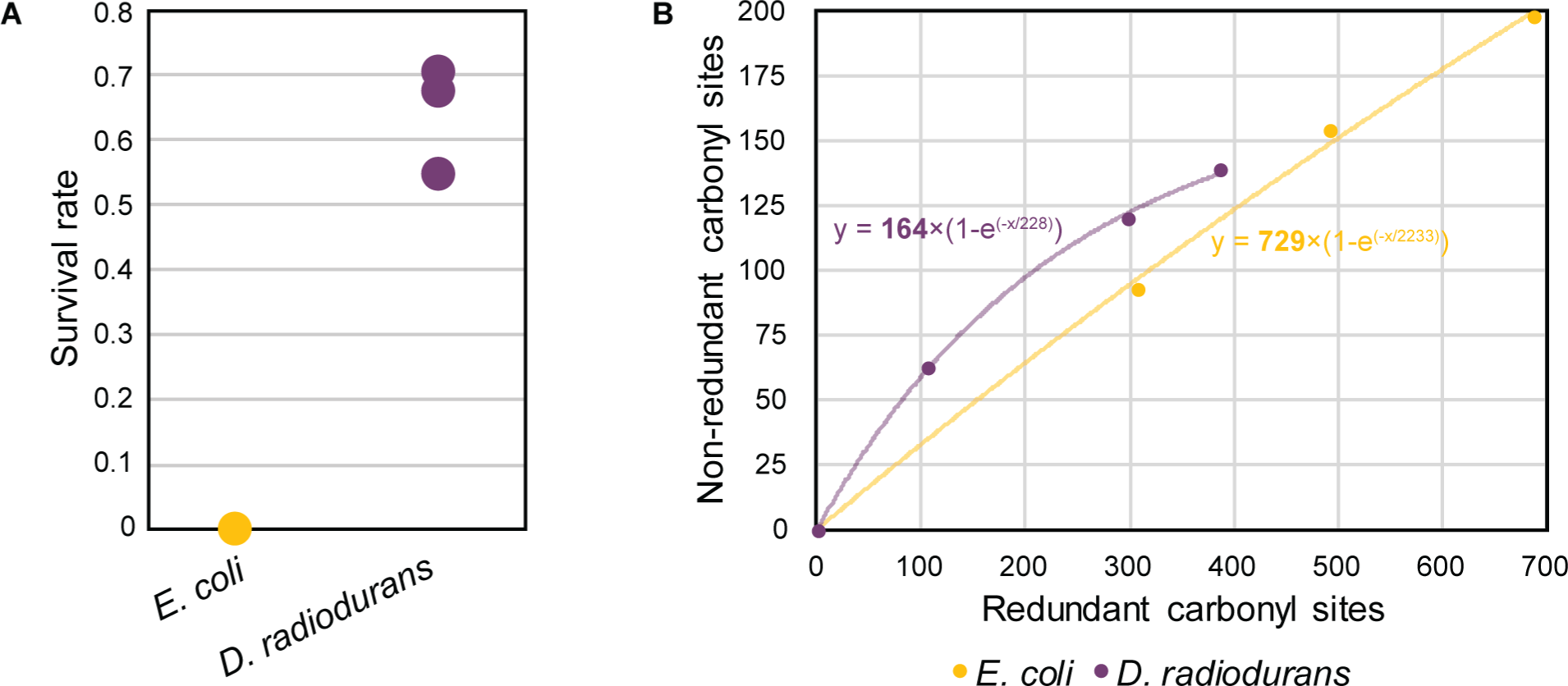
Survival and carbonyl site sampling limits for proteomic experiments, related to Fig. 2 and 3. (A) Survival rates (based on CFU counts) of irradiated *E. coli* and *D. radiodurans* corresponding to triplicate samples from which proteomic data was acquired. Absolutely no colonies were recovered from E. coli cultures that had been irradiated, even without diluting the samples before plating. (B) Carbonyl site measurement saturation curves for triplicate shotgun redox proteomic measurements in *E. coli* and *D. radiodurans*. Exponential saturation functions were fit by minimizing the sum of squared errors with the triplicate data points; the bolded term in each function is the estimated number of total non-redundant carbonyl sites in our samples.

**Fig. S3.**
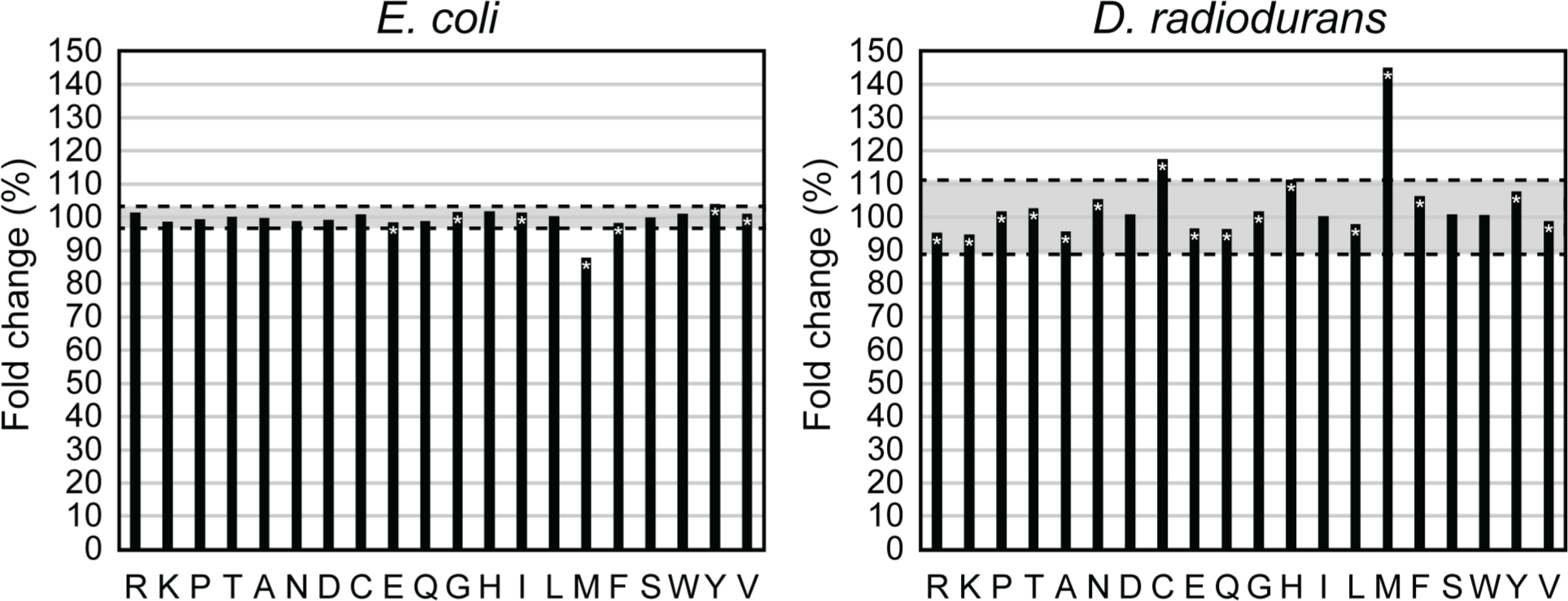
Canonical amino acid prevalence change following irradiation of *E. coli* (left) and *D. radiodurans* (right), related to Fig. 3. Shaded region indicates ± 1 standard deviation from the mean across all amino acids. * = z-test of proportions p-value < 0.01.

**Fig. S4.**
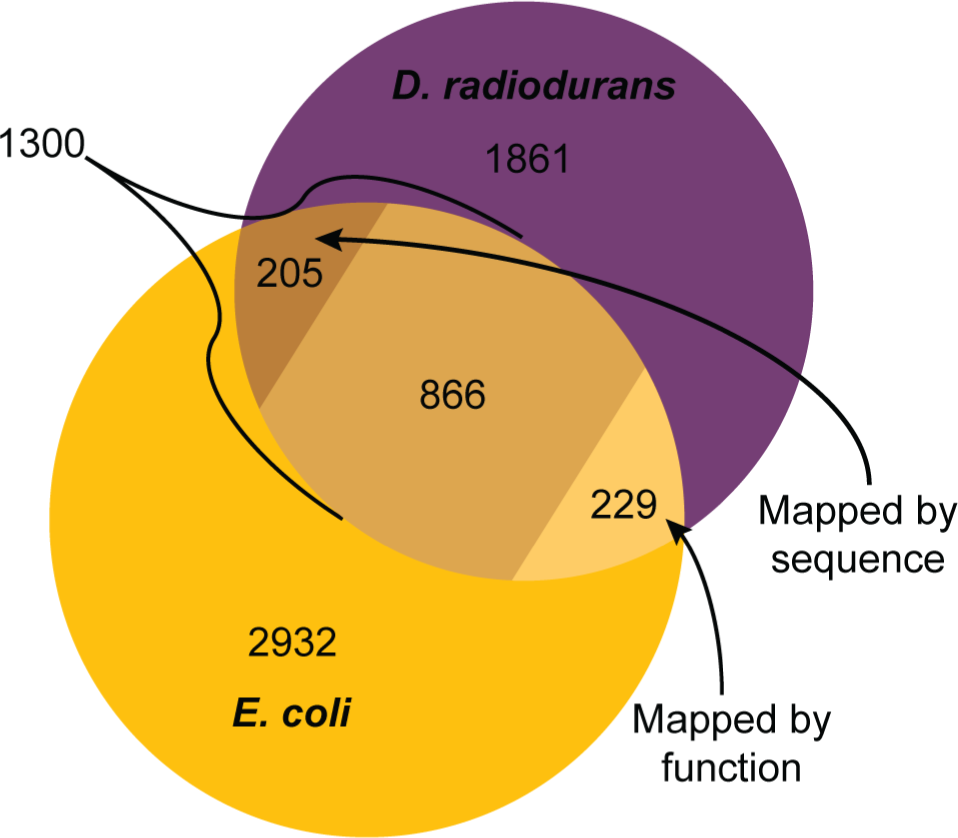
Orthologs mapped between *E. coli* and *D. radiodurans*, related to Fig. 7. 1300 orthologous pairs were mapped by manual curation between species aided by a combination of sequence alignment by reciprocal BLAST and functional annotation comparison using RAST. These orthologs were used to perform comparisons depicted in Fig. 7.

**Fig. S5.**
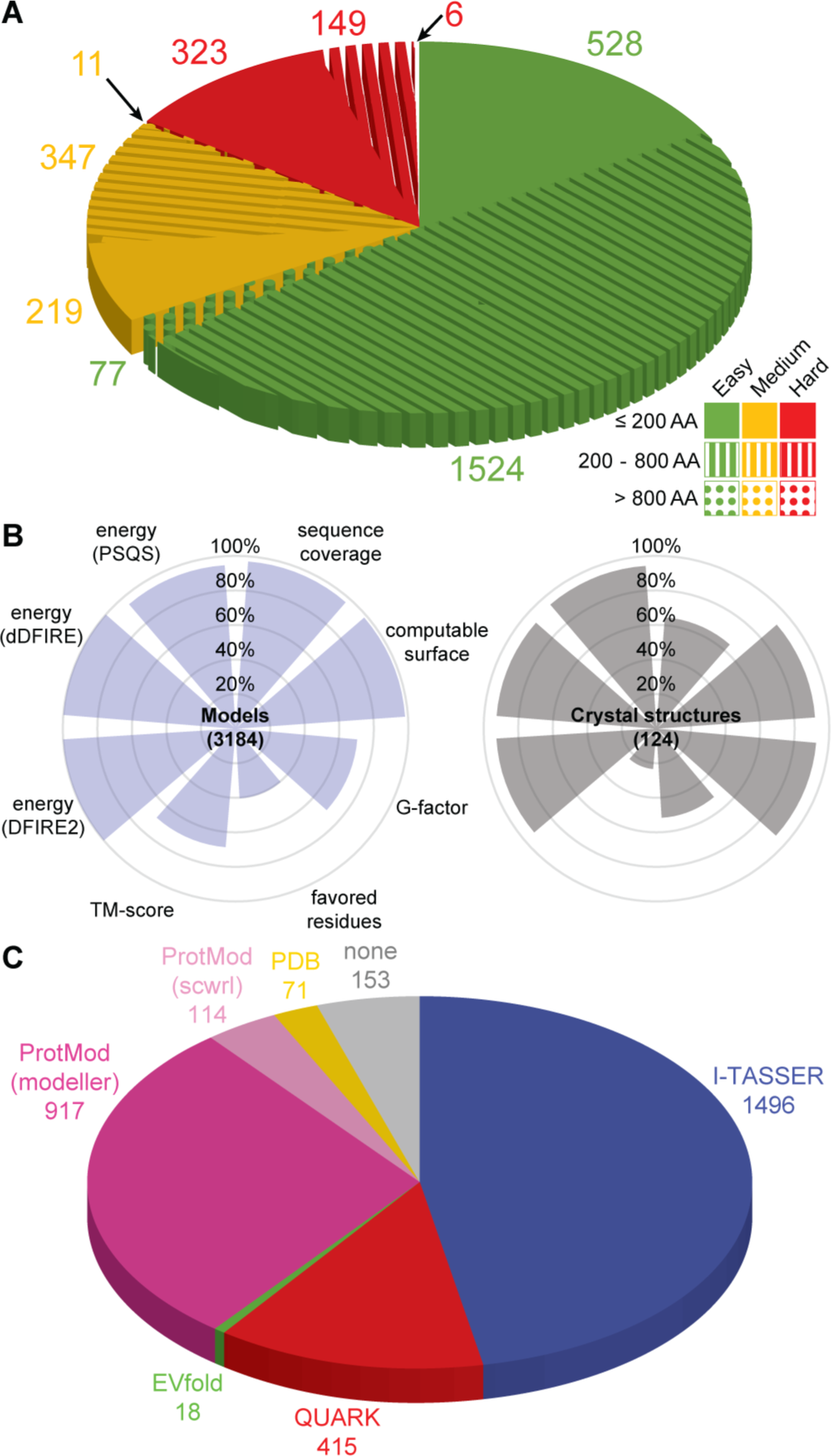
*D. radiodurans* proteome structure modeling, related to Fig. 4, 5, and 6. (A) Distribution of D. radiodurans proteins by difficulty of template-based homology modeling and size regimes relevant for determining structure modeling algorithm applicability. Easy signifies ≥10 high-confidence homologous templates available. Medium signifies ≥1 high-confidence homologous template available. Hard signifies no high-confidence homologous templates available. Proteins ≤ 200 residues long are amenable to ab initio folding. Proteins ≤ 800 residues long are amenable to homology modeling. (B) Structure quality evaluation criteria and percentage of *D. radiodurans* protein structures that satisfy published criteria thresholds. Blue plot represents best-representative models for *D. radiodurans* proteins. Gray plot represents best available crystal structures from the PDB for *D. radiodurans* proteins. (C) Distribution of methods used to derive best-representative protein structures for *D. radiodurans*. “None” indicates the proteins for which no PDB structure exists and no modeling method is applicable.

**Fig. S6.**
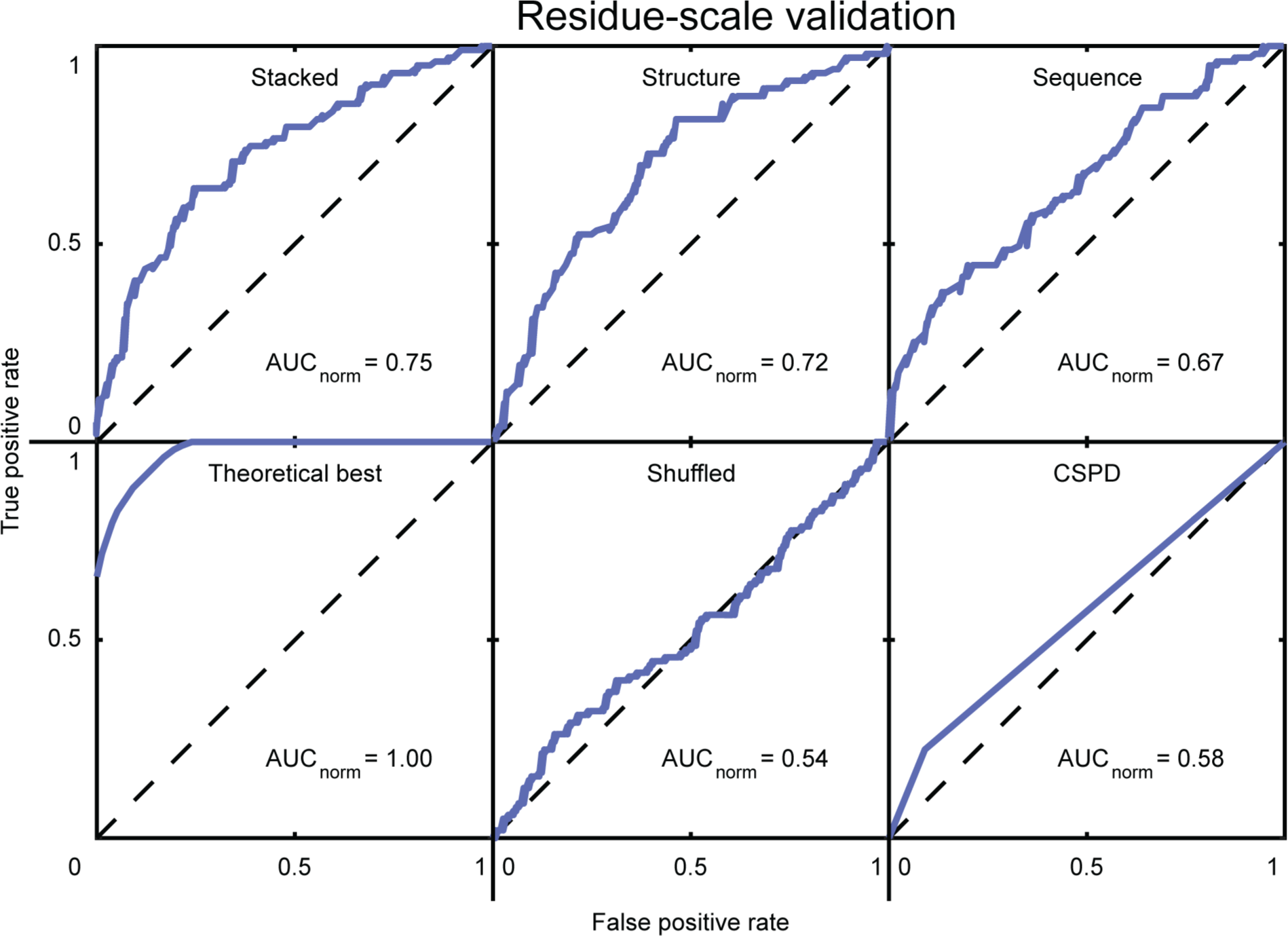
Residue-scale validation of metal-catalyzed oxidation (MCO) predictor, related to Fig. 5A. We developed an MCO predictor using the algorithm presented in this study but trained on previously published MCO data from *E. coli*, the same dataset used to develop CSPD. Receiver operating characteristic (ROC) curves for MCO site predictors derived by leave-1-out validation. The dashed black line at y=x corresponds to performance expected by chance. Top left = final predictor trained by stacking structure- and sequence-based models. Top middle = predictor trained only on structure-based features. Top right = predictor trained only on sequence-based features. Bottom left = theoretical maximum predictive power for a probability estimator (AUC = 0.97). Bottom middle = same algorithm as used for final predictor but with all features shuffled beforehand. Bottom right = CSPD model developed using MCO site data from *E. coli*.

**Fig. S7.**
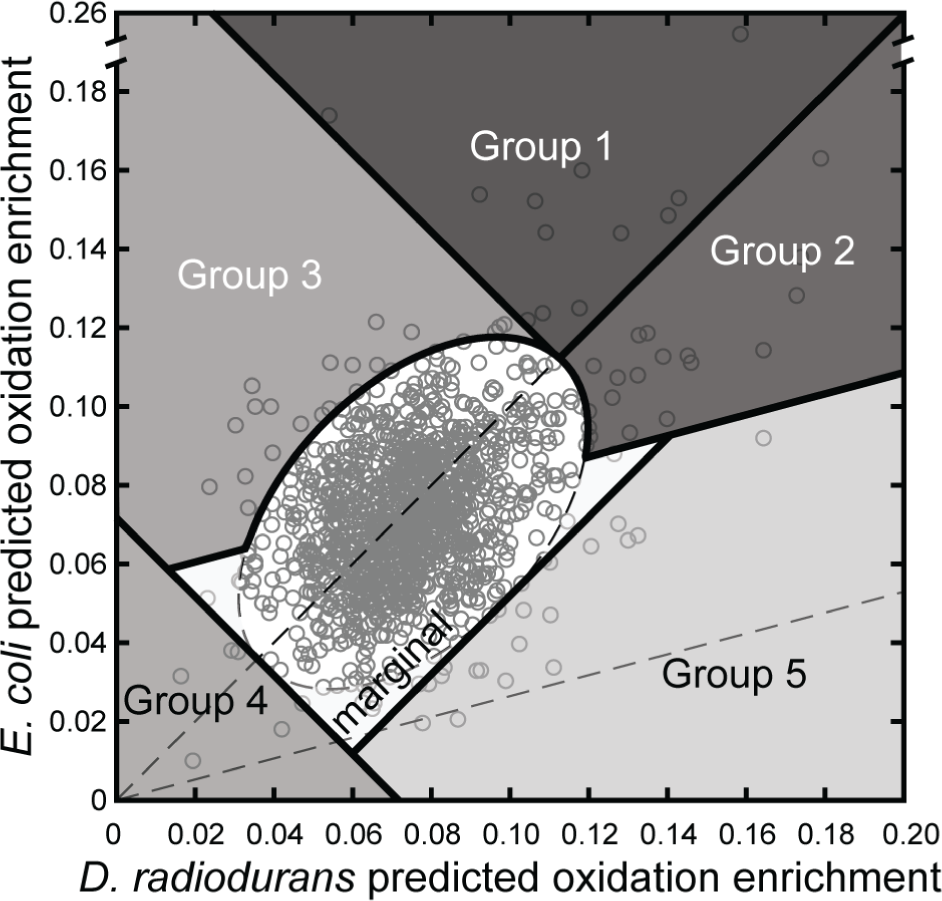
Predicted outliers grouped by comparative intrinsic and extrinsic vulnerability to oxidation in D. radiodurans and *E. coli*, related to Fig. 7. Group 1: Proteins predicted significantly more intrinsically prone to oxidation than the rest of the proteome in both species (>3σ distance from mean and above a line perpendicular to y=x that is tangent to the ellipse upper vertex) but more intrinsically (left of y=x) and extrinsically more protected in *D. radiodurans* (orthogonal distance from y=x/3.78 is significantly greater than the mean; p-value < 0.01). Group 2: Proteins predicted significantly more intrinsically prone to oxidation than the rest of the proteome in both species, with similar intrinsic vulnerability between species (<3σ distance from y=x but not in Group1) but more extrinsically protected in *D. radiodurans*. Group 3: Proteins predicted significantly more intrinsically (>3σ distance above y=x) or extrinsically protected against oxidation in *D. radiodurans* but not in Groups 1 or 2. Group 4: Proteins predicted significantly more intrinsically robust to oxidation than the rest of the proteome in both species (>3σ distance from mean and below a line perpendicular to y=x that is tangent to the ellipse lower vertex). Group 5: Proteins predicted significantly more intrinsically susceptible to oxidation in *D. radiodurans* than in *E. coli* (>3σ distance below y=x) and not significantly more extrinsically protected than the rest of the proteome (orthogonal distance from y=x/3.78 is not significantly greater than the mean). Marginal: Proteins not predicted to have any significant intrinsic or extrinsic protection in *D. radiodurans* or *E. coli*.

Table S1. (separate file)

All peptides detected in redox proteomic experiments. *D. radiodurans* and *E. coli* results are on separate worksheets. With exception of organism, condition, replicate, and carbonylation columns, all columns are as directly output by ProteinPilot.

Table S2. (separate file)

All quantitative proteomics data. *D. radiodurans* and *E. coli* results are on separate worksheets.

Table S3. (separate file)

*D. radiodurans* protein structure summary with quality evaluation metrics.

**Table S4.** Structure derivation method applicability.

| | Easy<br>($\geq 10$ templates)<sup>a</sup> | | | | | Medium<br>(1 - 10 templates)<sup>a</sup> | | | | | Hard<br>(0 templates)<sup>a</sup> | | |
| --- | --- | --- | --- | --- | --- | --- | --- | --- | --- | --- | --- | --- | --- |
| $\leq 200$ AA | P | I | M | Q | E | P | I | M | Q | E | P | Q | E |
| 200 - 800 AA | P | I | M |  | E<sup>b</sup> | P | I | M |  | E<sup>b</sup> | P |  | E<sup>b</sup> |
| 800 - 1500 AA | P | I | M |  |  | P | I | M |  |  | P |  |  |
| 1500 - 2000 AA | P |  | M |  |  | P |  | M |  |  | P |  |  |
| > 2000 AA | P |  |  |  |  | P |  |  |  |  | P |  |  |

P = PDB (experimental)
I = I-TASSER (local homology)
M = ProtMod (global homology)
Q = QUARK (ab initio folding)
E = EVfold (evolutionarily-constrained ab initio folding)
<sup>a</sup>High confidence templates found by LOMETS
<sup>b</sup>Transmembrane proteins

**Table S5.** Protein structure quality evaluation metrics.

| Metric | Threshold | Tool |
| --- | --- | --- |
| Sequence coverage | > 90%<sup>a</sup> | (align to target) |
| Computable surface | TRUE<sup>a</sup> | UCSF Chimera |
| Energy scores | < 0 | PSQS, dDFIRE, DFIRE2 |
| TM-score (estimated) | > 0.5 | ResQ |
| Ramachandran favored residues | > 90% | PROCHECK |
| G-factor overall conformation | > -1 | PROCHECK |

<sup>a</sup>Absolute requirement

**Table S5.**
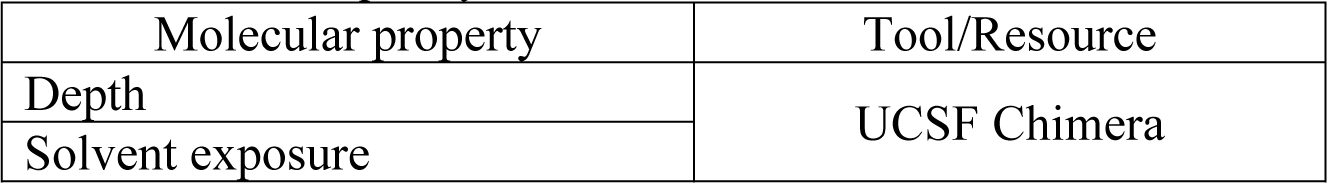
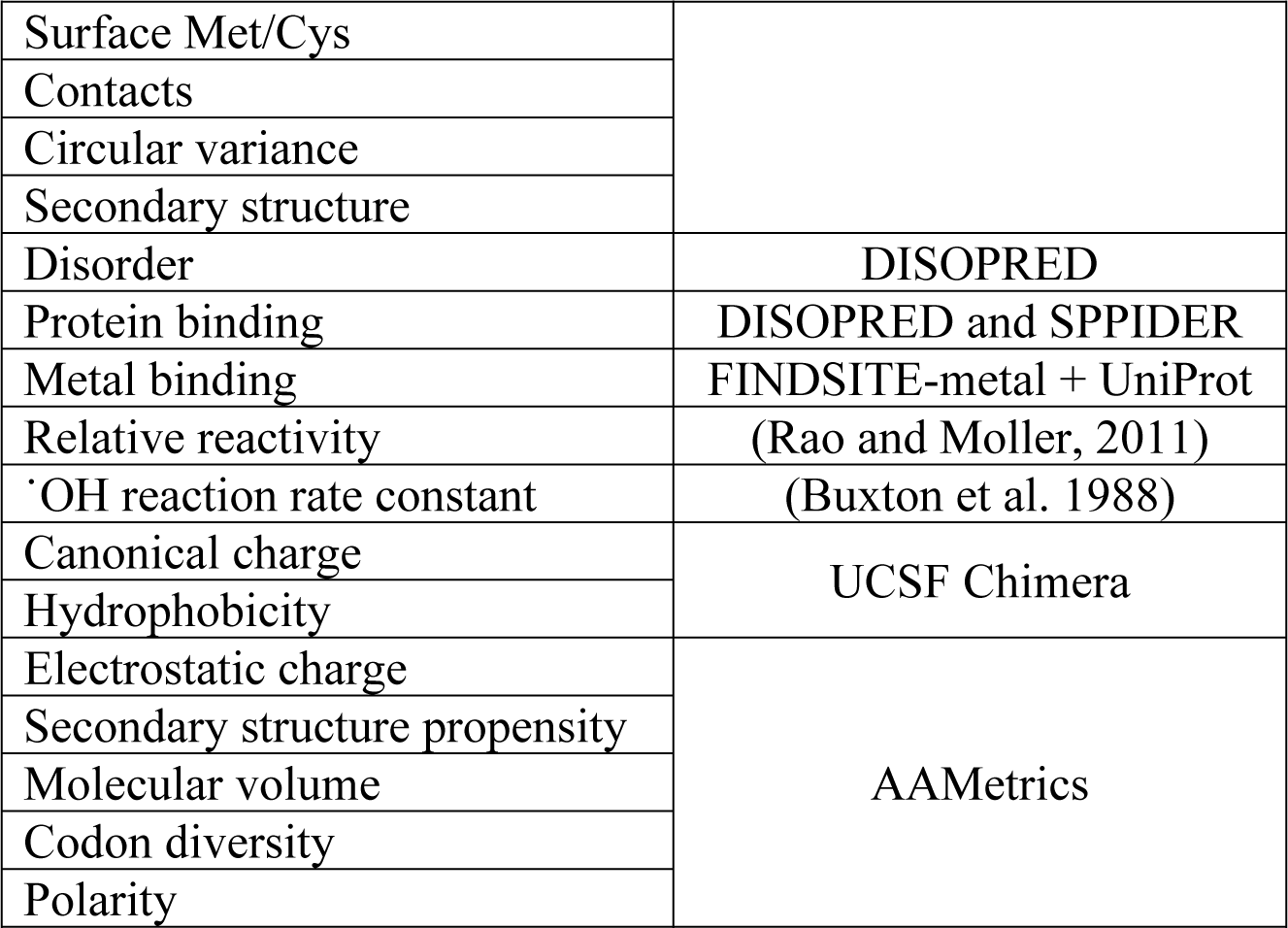
Protein structure quality evaluation metrics.

Table S7. (separate file)

Carbonyl site training data with all features.

Table S8. (separate file)

*E. coli* carbonyl site predictions proteome-wide. Only abbreviated feature data are included, structure-based features included in the final predictor.

Table S9. (separate file)

*D. radiodurans* carbonyl site predictions proteome-wide. Only abbreviated feature data are included, structure-based features included in the final predictor.

Table S10. (separate file)

Protein vulnerability to oxidation predictions and experimental data summary. *D. radiodurans*,

*E. coli*, and interspecies comparison results are on separate worksheets.

File S1. (separate file)

Irradiator rack body STL file.

File S2. (separate file)

Irradiator rack lid STL file.

File S3. (separate file)

ProteinPilot data dictionary.

File S4. (separate file)

ProteinPilot parameter translation.

